# Chemogenetic timestamping for the precise tracing of cell history into protein assemblies

**DOI:** 10.64898/2026.07.10.737712

**Authors:** Lina El Hajji, Arnaud Gautier

## Abstract

Self-assembling protein fibers enable to record events in single cells, bypassing the need for long-term time-lapse imaging. Fluorescent marks introduced within the growing fiber at user-defined times provide timestamps, giving access to the temporal dynamics of the recorded event. Here, we introduce CATCHFiber, a single-color timestamping strategy for tracing cellular events with high temporal resolution into self-assembling protein fibers. Relying on chemically-induced dimerization to precisely and rapidly control the incorporation of fluorescent proteins into the fiber, CATCHFiber allows the introduction of short 30-min spaced timestamps, significantly increasing the precision of event timings compared to existing methods. This increase in temporal resolution expands the use of fiber-based recorders beyond transcriptional activity, allowing to trace the kinetics of faster processes such as protein degradation, protein neosynthesis and kinase activity, and to determine the timing of cell cycle steps.

---

The intricate biological processes orchestrating global cell function span over a wide range of timescales. Fluorescence microscopy allows real-time tracking of cellular events using time-lapse imaging. Through adjusting the acquisition frame-rate and choosing adequate fluorescent reporters, time-lapse-imaging allows to capture both fast-occurring and long-term processes. However, long-term replicated measurements of large cell populations involve time-consuming experiments, and require the handling and storage of large datasets.

Recently, large-scale single-cell analysis of biological processes was achieved by recording cellular events into protein fibers growing within cells^1^. In this approach, an event such as gene activation leads to the incorporation of a fluorescent reporter into the fiber, leaving a permanent mark. The precise timing of the event is obtained through embedding user-defined timestamps along the fiber (**Figure 1a**). Recording events by spatially encoding the temporal information suppresses the need for labor-intensive and time-consuming time-lapse microscopy. Post-hoc snapshots of fibers give access to the temporal information and kinetics (**Figure 1a)**, enabling highly parallelized, large-scale acquisitions, with information at the single cell-level.

**Figure 1.**
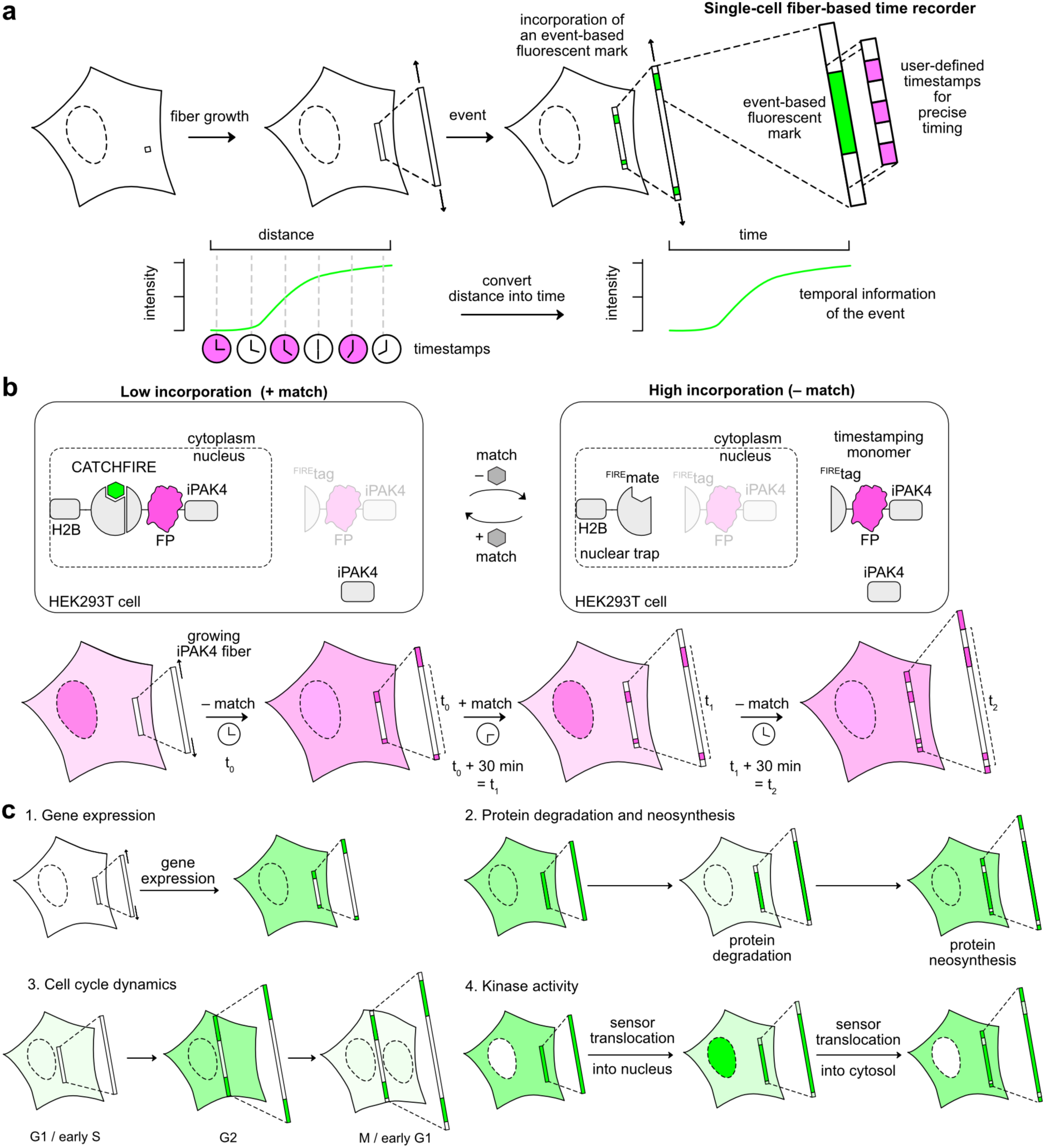
CATCHFiber – a chemogenetic strategy for tracing cellular events into protein fibers. **a** General principle of fiber-based recorders. An event of interest leads to the incorporation of a fluorescent mark within a growing protein-based fiber. Precise timing is obtained through simultaneous incorporation of user defined timestamps. These timestamps enable to convert the intensity profile of the event-based fluorescent mark along the fiber into a temporal information. **b** iPAK4 monomers can form linear fibers into cells. Timestamping is achieved through the controlled incorporation of a fluorescently labeled iPAK4 (timestamping monomer) using CATCHFIRE, a chemically-induced dimerization tool. The timestamping monomer contains a ^FIRE^tag peptide, which interact in a reversible manner with a ^FIRE^mate domain in presence of a match molecule. Addressing ^FIRE^mate in the nucleus (through fusion to H2B) enables to modulate the cytosolic pool of timestamping monomers in a match-dependent manner. Repetitive match washout and addition enables to create timestamps into growing iPAK4 fibers. **c** Summary of the different processes recorded and analyzed using CATCHFiber-based timestamping in this study.

Two strategies were developed for recording temporal information into fibers. The first one – expression recording islands (XRI) – relies on the isoaspartyl dipeptidase 1POK (E239Y) protein, which forms intracellular flexible slow-growing fibers well suited for long-term recording of gene activity in mammalian cells^2^. The second one uses iPAK4 – a fusion of the catalytic domain of PAK4 kinase and the iBox domain of its inhibitor Inka1^3^ – which forms rigid intracellular fibers with limited toxicity in mammalian cells^4^. Multicolor timestamping was achieved through incorporation of HaloTag-iPAK4 monomers labeled at user-defined timepoints with different fluorescent dyes, allowing the recording of gene activation in mammalian cells and neurons^3^. Computational engineering enabled the development of CytoTape, an improved version of the XRI system, allowing weeks-long multiplexed recordings of transcriptional activity^5^. The use of the iPAK4 ticker-tape system was also extended for long-term multiplexed parallelized recording through inducible expression of fluorescent proteins (FP)^6^.

The growing interest for fiber-based recorders revealed the need for refined and improved timestamping strategies to expand their scope of applicability. Current timestamping methods are limited by their temporal resolution. HaloTag timestamps can be spaced every two hours at best, because of the long half-life (4.5 h) of HaloTag^3^. Similarly, approaches relying on conditional expression of FPs are limited to three-hours-timestamps because of the delayed response of gene activation^6^. In addition, in both strategies, the use of multiple colors for timestamping monopolizes several imaging channels, complicating multiplexed recordings. Here, we present CATCHFiber – Chemically-Assisted Tracing of Cell History into Fibers –, a single-color timestamping strategy with enhanced temporal resolution. CATCHFiber enables to incorporate narrowly-spaced timestamps within iPAK4 fibers through user-controlled trafficking of fluorescent timestamp monomers in and out of the cell nucleus using reversible chemically-induced dimerization (CID) (**Figure 1b**). This approach conditions the incorporation of the timestamping monomers into the fiber to the rapid addition or removal of a small molecule, allowing user-defined control. The kinetics of CATCHFiber enable the introduction of timestamps every 30 minutes, resulting in increased temporal resolution in comparison to existing strategies. We show that CATCHFiber opens new perspectives for the use of fiber-based cellular recorders beyond the recording of transcriptional events, enabling accurate recording of fast cellular events such as ubiquitination-mediated degradation of proteins, protein neosynthesis and the activity of kinases involved in cell signaling, and dynamic cellular processes such as the cell cycle (**Figure 1c**).

## RESULTS

### Timestamping with CATCHFIRE

The original iPAK4-based ticker tape works by combining a large excess of iPAK4 monomers – driving fiber formation – with a smaller fraction of fluorescently-labeled HaloTag-iPAK4 fusion for timestamping. Here, we propose that an alternative way of timestamping is to modulate the available cytosolic pool of timestamp monomers through repetitive and user-controlled cycles of sequestration and release in and out the cell nucleus. To do so, we relied on the chemically assisted tethering of chimera by fluorogenic-induced recognition (CATCHFIRE) approach, a fluorogenic chemicallyinduced dimerization (CID) tool composed of two protein fragments called ^FIRE^mate and ^FIRE^tag forming a fluorescent ternary assembly in the presence of a small fluorogenic dimerizer called match. The ternary fluorescent assembly forms rapidly and can be efficiently dissociated by washing away the match molecule^7^.

CATCHFIRE was previously shown to allow passive nuclear import of small ^FIRE^tagged cytosolic proteins through match-induced interaction with a nuclear ^FIRE^mate anchor via a sink effect^7^. We thus reasoned that fusing ^FIRE^tag to timestamp monomers containing fluorescent proteins (FP) would allow to control their intracellular localization through match-dependent interaction with a co-expressed H2B-^FIRE^mate nuclear trap, enabling the incorporation of closely-spaced fiducial fluorescent timestamps within iPAK4 fibers (**Figure 1b**).

Co-expression of iPAK4 (90%) with ^FIRE^tag-FP-iPAK4 (FP = EGFP, mCherry or emiRFP670) fusions (10 %) generated fluorescent fibers similar to the ones generated using FP-iPAK4 (**Supplementary Figures 1, 2 and 3**), showing that ^FIRE^tag did not affect neither fiber formation and growth nor incorporation of the FP into the fiber. In addition, two FP could be simultaneously incorporated into the fibers, simulating situations where an FP for timestamping and one for event recording are used (**Supplementary Figures 4 and 5**).

**Figure 2.**
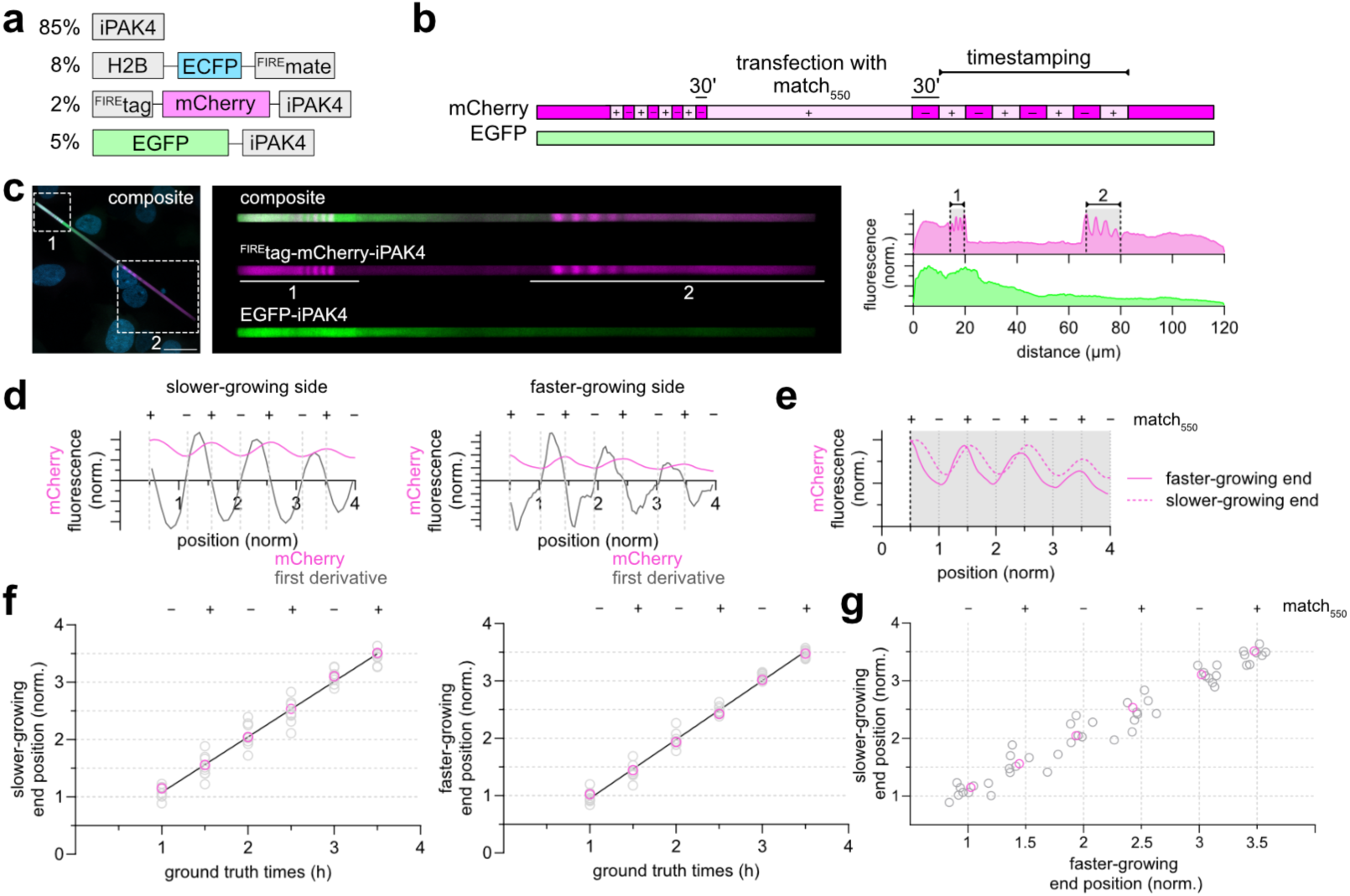
Evaluation of both iPAK4 fiber ends for CATCHFiber timestamping. **a** Plasmids used. **b-g** HEK293T cells expressing CATCHFiber_red_ were subjected to the timestamping sequence match_550_[30− / 30+]_4h_. **c** A representative micrograph as well as the fluorescence intensity plot along the fiber is shown. Scale bar 20 µm. The timestamping period is shown as a grey highlight in the full fluorescence intensity plot. A straightened image of the full fiber is also shown. The dashed black lines positions were normalized to 0.5 and 4. **d** Graphs showing the fluorescence intensity (color) and the first derivative (gray) as a function of the normalized position for the slower growing end (1, left) and for the faster growing end (2, right). **e** Superimposition of the normalized fluorescence intensity from the two fiber ends as a function of the normalized position. The results are representative of n = 9 fibers from four replicates. **f** Positions of the +/- match_550_ transitions (grey) – identified as the cancellation points of the first derivative of the intensity profile – are plotted vs the ground truth times of match_550_ washes and additions for the slower-growing end (left) and the faster growing end (right). In pink are shown the transitions identified for the fiber displayed in (**c**). **g** Positions of the match transitions for the slower growing end of the fiber against the corresponding position for the faster growing end (grey). In pink are shown the positions for the fiber displayed in (**c**).

**Figure 3.**
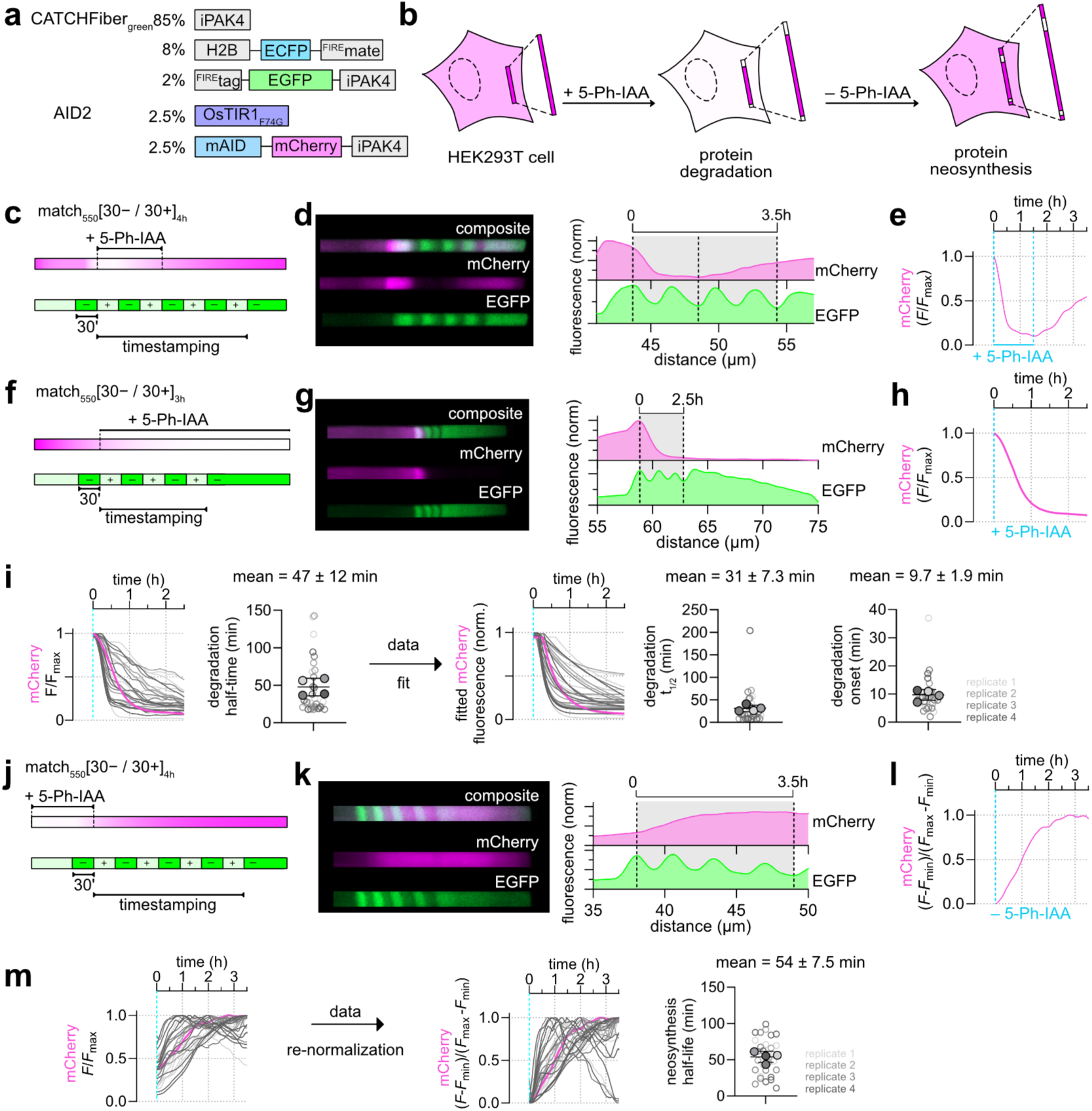
CATCHFiber enables the recording of proteasomal protein degradation and neosynthesis. **a** Plasmids used. **b** Proteasomal protein degradation was induced by addition of 5-Phenyl-indole-3-acetic acid (5-Ph-IAA, 1 µM), and neosynthesis was enabled through washing out 5-Ph-IAA. **c-e** Evaluation of the reversibility of the AID2 system using CATCHFiber. HEK293T cells expressing CATCHFiber_green_ and AID2 were subjected to the timestamping sequence match_550_ [30− / 30+]_4h_. A zoomed-in image of a representative fiber end and the corresponding intensity plot are shown in (**d**). The dashed black lines positions were normalized to 0 and 3.5 h. Degradation was induced through addition of 5-Ph-IAA along with the first match_550_ addition, and neosynthesis was induced through washing out 5-Ph-IAA 1.5 h later. **e** Temporal evolution of mCherry intensity after temporal rescaling (see methods and **Supplementary Text 1** for detailed calculations). The blue dashed lines indicate the duration of incubation in the presence of 5-Ph-IAA (1 µM). The results are representative of n = 33 fibers from two replicates. **f-i** Evaluation of the AID2-induced protein degradation dynamics. HEK293T cells expressing CATCHFiber_green_ and AID2 were subjected to the timestamping sequence match_550_ [30− / 30+]_3h_. Degradation was induced through addition of 5-Ph-IAA along with the first match_550_ addition. A zoomed-in image of a representative fiber end and the corresponding intensity plot are shown in (**g**). The dashed black lines positions were normalized to 0 and 2.5 h. **h** Temporal evolution of mCherry intensity after temporal rescaling (see methods and **Supplementary Text 1** for detailed calculations). The blue dashed line indicates the addition of 5-Ph-IAA (1 µM). The results are representative of n = 31 fibers from four replicates. **i** Temporal evolution of normalized mCherry intensity from all 31 fibers imaged in this experiment after temporal rescaling. Each curve is color-coded according to the replicate the fiber comes from. In pink is the temporal evolution of the fiber shown in **g.** For each curve, the time position corresponding to 50% of the maximal mCherry signal is reported as the degradation half-time. In this graph, each fiber is color-coded according to the biological replicate it came from. The solid circles correspond to the mean of each biological replicate. The black line represents the mean ± SD of the four replicates (n = 31 fibers). The curves are then fitted using a one-phase decay model with an initial plateau, and for each fitted curve, are extracted and plotted the degradation half-life and onset. In each plot, each fiber is color-coded according to the biological replicate it came from. The solid circles correspond to the mean of each biological replicate. The black line represents the mean ± SD of the four replicates (n = 31 fibers). **j-m** Evaluation of the neosynthesis dynamics following 5-Ph-IAA washout. HEK293T cells expressing CATCHFiber_green_ and AID2 pretreated with 5-Ph-IAA were subjected to the timestamping sequence match_550_[30− / 30+]_4h_. Neosynthesis was induced through washing out of 5-Ph-IAA along with the first match_550_ addition A zoomed-in image of a representative fiber and the corresponding intensity plot are shown in (**k**). The dashed black lines positions were normalized to 0 and 3.5 h. **l** Temporal evolution of mCherry intensity after temporal rescaling (see methods and **Supplementary Text 1** for detailed calculations). The blue dashed line indicates the washing out of 5-Ph-IAA (1 µM). The results are representative of n = 29 fibers from four replicates. **m** Temporal evolution of normalized mCherry intensity from all 29 fibers imaged in this experiment (grey, pink) after temporal rescaling. Each curve is color-coded according to the replicate the fiber comes from. In pink is the temporal evolution of the fiber shown in **k.** For each curve, the time position corresponding to 50% of the maximal mCherry signal is reported next as the neosynthesis half-life. In this graph, each fiber is color-coded according to the biological replicate it came from. The solid circles correspond to the mean of each biological replicate. The black line represents the mean ± SD of the four replicates (n = 29 fibers).

**Figure 4.**
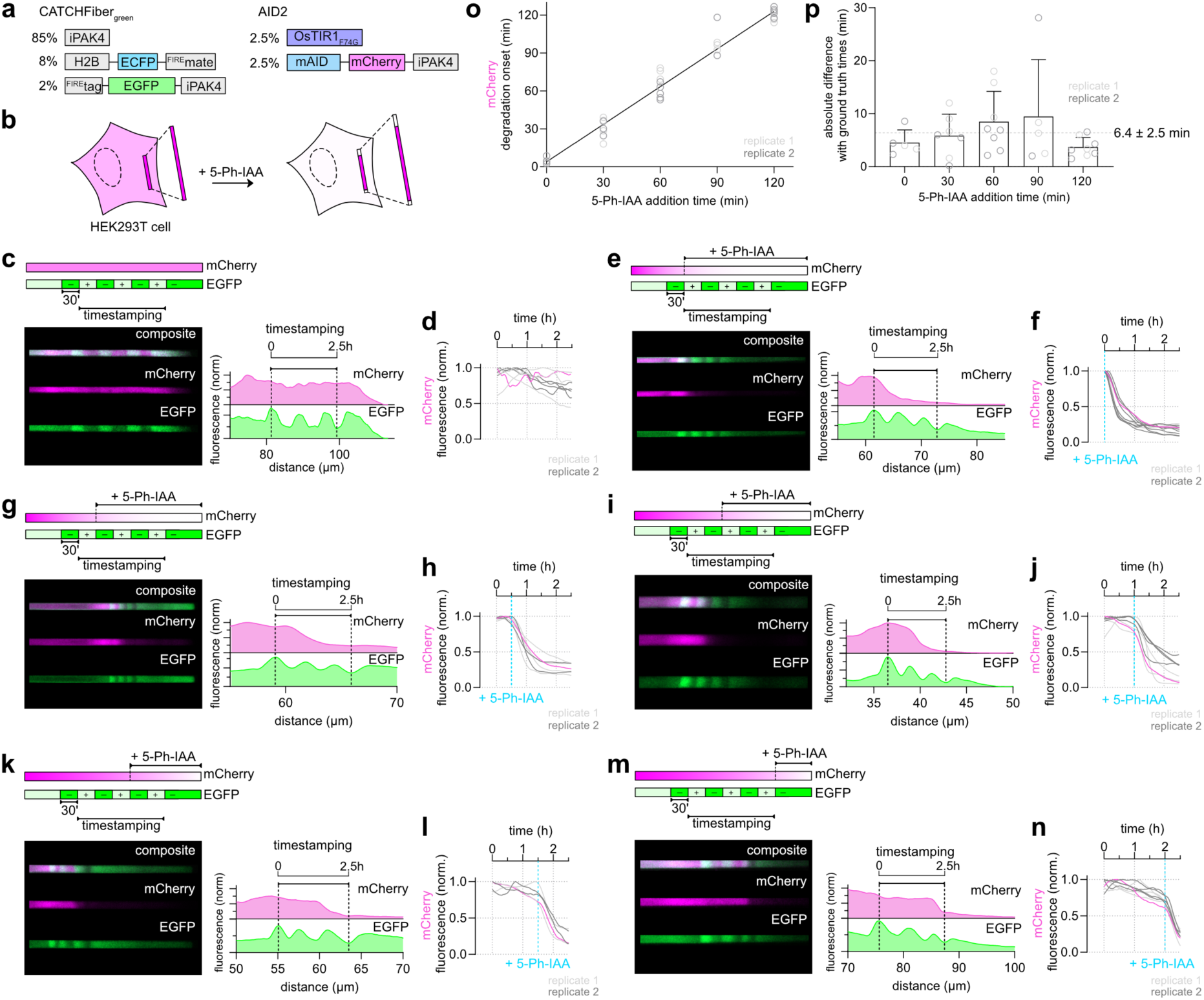
CATCHFiber reports on the timing of protein degradation onset. **a** Plasmids used. **b** Proteasomal degradation was induced through the addition of 5-Ph-IAA (1 µM). HEK293T cells expressing CATCHFiber_green_ along with AID2 were subjected to the timestamping sequence match_550_ [30− / 30+]_3h_ and subjected to no (**c**,**d**) or 1 µM of 5-Ph-IAA (**e-n**) added at t = 0 (**e,f**), t = 30 min (**g,h**), t = 1 h (**i,j**), t = 1.5 h (**k,l**) and t = 2 h (**m, n**) after timestamping start. For each condition are shown a zoomed-in image of a representative fiber and the corresponding intensity plot (**c,e,g,i,k,m**). Representative results from n = 8 fibers (**c, d**), n = 10 fibers (**e,f**), n = 7 fibers (**g,h**), n = 8 fibers (**i,j**), n = 5 fibers (**k,l**), and n = 8 fibers (**m,n**) from two replicates. Temporal evolution of normalized mCherry intensity from all fibers after temporal rescaling are also shown (**d,f,h,j,l,n**). Each curve (grey) is color-coded according to the replicate the fiber comes from. In pink is the temporal evolution of the representative fiber shown left. **o** Degradation onset for the curves shown in **f, h, j, l, n**, identified as local minima in the second derivative of the intensity plot. 5 curves were omitted from **f**, as the degradation happened without measurable onset. Otherwise, all curves were considered for analysis and all data points were shown. **p** Absolute difference between degradation onsets and ground truth times of addition of 5-Ph-IAA.

**Figure 5.**
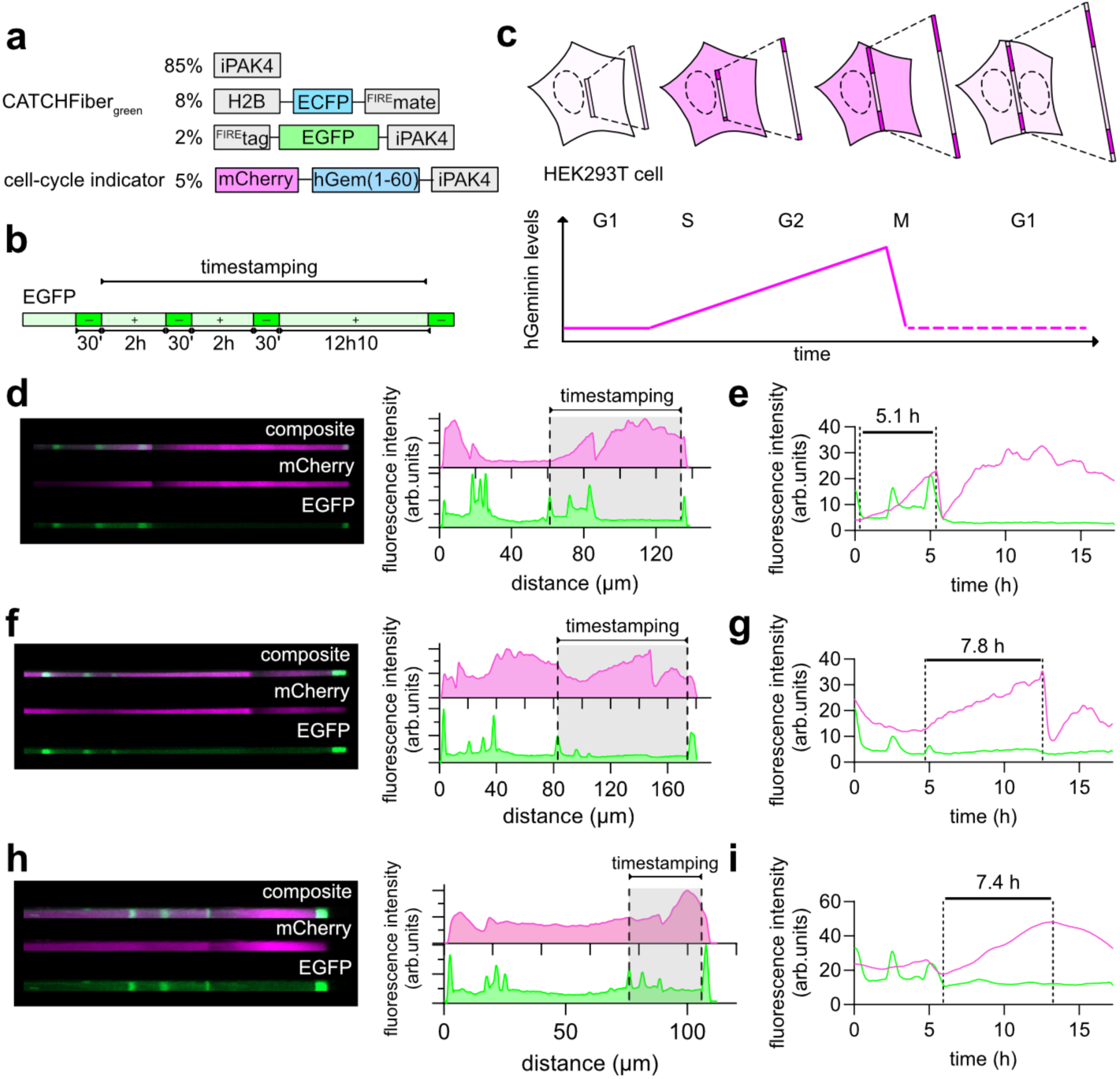
CATCHFiber enables recording the cell-cycle dynamics. **a** Plasmids used. **b** HEK293T cells expressing CATCHFiber_green_ along with a cell-cycle indicator were submitted to the indicated timestamping sequence. **c** When cells progress through the cell cycle, the expressed iPAK4 fibers show varying levels of mCherry, correlating with the cell-cycle dependent evolution of human geminin (hGem) levels. A zoomed-in image of three representative fiber ends and the corresponding intensity plot of the full fibers are shown (**d, f, h**). Temporal evolution of normalized mCherry intensity after temporal rescaling is shown (**e, g, i**). The durations of the S-G2 phases, identified as periods of increase in mCherry levels are indicated. Representative results from n = 19 fibers from four replicates.

User-defined control of the intracellular localization of timestamp monomers was demonstrated transfecting HEK293T cells with plasmids encoding H2B-ECFP-^FIRE^mate (80%) and ^FIRE^tag-mCherry-iPAK4 (20%) (**Supplementary Figure 6a,b**). Time-lapse imaging revealed a homogeneous localization of ^FIRE^tag-mCherry-iPAK4 in the cell in the absence of match_550_. Addition of 10 µM of match_550_ led to the association with ^FIRE^mate and ^FIRE^tag as demonstrated by the appearance of a green fluorescent signal in the nucleus, a decrease of the mCherry cytosol/nucleus ratio and an increase of nuclear mCherry (**Supplementary Figure 6c-e**).

We leveraged this property to precisely place fiducial patterns within iPAK4 fibers. We anticipated that user-controlled match addition would lead a dark mark on the growing fiber through trapping of the timestamping monomers in the nucleus, while conversely, match washing would result in a bright mark because of their release in the cytosol and incorporation in the fiber. Here, the duration of the +/– match pulses would define the timestamps spacing. To test this, we transfected HEK293T cells with plasmids encoding iPAK4 (90%), H2B-ECFP-^FIRE^mate (8%), and ^FIRE^tag-EGFP-iPAK4 (2%) (**Supplementary Figure 7a-j**). Cells were grown overnight in the presence of 10 µM match_550_ to trap the newly synthesized ^FIRE^tag-EGFP-iPAK4 in the nucleus. The following day cells were subjected to either 2 h, 1 h or 30 min alternating incubations without and with match_550_ for a total duration of 4h, and then maintained without match_550_ prior to imaging. To improve readability, we introduce the following terminology: match_550_[X– / Y+]_Zh_ corresponds to a sequence of Z hours, consisting of alternating X minutes of – match_550_ incubation and Y minutes of + match_550_ incubation. The fibers from match_550_[30− / 30+]_4h_ (**Supplementary Figure 7b-d**), match_550_[60− / 60+]_4h_ (**Supplementary Figure 7e-g**) and match_550_[120− / 120+]_4h_ (**Supplementary Figure 7h-j**) sequences displayed low fluorescence at their center, and alternating bright and dark bands on both edges of the fibers in the green channel, matching the alternating absence and presence of match_550_ (**Supplementary Figure 7d,g,j**). Similar results were obtained using ^FIRE^tag-mCherry-iPAK4 as timestamp monomer (**Supplementary Figure 7k-t**).

To precisely analyze timestamping, we developed a dedicated analysis pipeline (see **Materials and Methods**, **Annex Figures** and **Supplementary Text 1** for detailed analysis explanation). Briefly, the maxima and minima of the fluorescence intensity plot curve were identified looking at the cancellation points of the first derivative, enabling us to precisely position the match additions and washouts. These points were in average equally spaced for the match_550_[30− / 30+]_4h_ pulse sequence (**Supplementary Figure 7d,n**), in agreement with a linear growth of the fiber in average. Similar behavior was observed for the pulse sequences match_550_[60− / 60+]_4h_ and match_550_[120− / 120+]_4h_ (**Supplementary Figure 7g,j,q,t**), although we observed a first phase of slight decrease followed by a second sharper decrease in fluorescence intensity following match addition. We thus retained pulses of 30 minutes for timestamping for clearer timestamping positioning.

We named our new timestamping strategy CATCHFiber (Chemically-Assisted Tracing of Cell History into Fiber), and added the suffix green (FP = EGFP) or red (FP = mCherry) to specify the timestamp color. The ability of CATCHFiber to impart more closely-spaced fiducial timestamps (30 minutes vs 2 hours for the HaloTag-based ticker tapes), in a single imaging channel, opens interesting perspectives for multiplexed recordings with high temporal resolution.

### Timestamping analysis

In agreement with iPAK4 fibers growing in both directions, we observed CATCHFiber timestamps present on both sides of iPAK4 fibers when using match_550_[30− / 30+]_4h_ pulse sequence (**Figure 2a-c**). Although one edge grows faster than the other, the two sides displayed almost superimposable fluorescent patterns after position normalization (**Figure 2d,e**), suggesting that both sides could be used indistinctively for event recording. For each side, the transitions corresponding to the wall clock times 1, 1.5, 2, 2.5, 3 and 3.5 h were positioned in average linearly along the fiber. However, analysis of individual fibers showed small cell-to-cell deviations to this average linear behavior (5.4 ± 1.6 min and 7.8 ± 1.8 min for the faster and slower growing end for all six transitions, respectively) because all fibers do not grow perfectly at constant rate during the 4 h period (**Figure 2f,g**). When recording events occurring over a short timescale, it appears crucial to place narrowly-spaced timestamps to increase temporal accuracy, since cells can undergo physiological changes affecting linear fiber growth. To maximize the number of analyzable fibers and benefit from maximal precision, we focused our analysis in the rest of the study on the faster growing side.

### Recording gene activation

To test the efficacy of CATCHFiber timestamping, we first recorded gene activation using a doxycycline-inducible system. We expressed rtTA3 and EGFP-iPAK4 under a tetracycline response element (TRE) promoter in HEK293T cells (**Supplementary Figures 8 and 9**). Timestamping was achieved using CATCHFiber_red_ together with a match_550_[30− / 60+]_4.5h_ pulse sequence (**Supplementary Figure 9a-c**). Gene expression was activated by adding doxycycline 30 min after timestamping initiation. The fibers displayed low EGFP fluorescence in the center and at the beginning of the timestamping period, and bright fluorescent edges, in agreement with the doxycycline induced expression of EGFP-iPAK4 and its subsequent incorporation into the fiber (**Supplementary Figure 9d,e**). Temporal rescaling using the timestamping information enabled to correct for potential deviations from the linear growth hypothesis (**Supplementary Figure 9f**, **Supplementary Text 1**, **Annex Figure 2**). Kinetics analysis showed that doxycycline-induced EGFP-iPAK4 expression occurs with an average onset of 70 ± 14 minutes (**Supplementary Figure 9g,h**), in agreement with the 1.2 h onset originally reported using HaloTag-based timestamping^3^.

### Recording protein degradation and neosynthesis

Protein-based recorders hold great potential for providing single-cell recordings beyond transcriptional activity. The increased temporal resolution of CATCHFiber timestamping enables to envision analysis of events with faster kinetics. As fast cellular events, we first considered proteasomal protein degradation and protein neosynthesis (i.e. translation only). We leveraged the auxin-inducible degron 2 (AID2) system, consisting of the Fbox Oryza sativa TIR1 F74G mutant (OsTIR1_F74G_) and mAID, a small 7 kDa truncation of the auxin-inducible degron from Arabidopsis IAA17^8^. Addition of 5-phenyl-indole-3-acetic acid (5-Ph-IAA) induces interaction between mAID and OsTIR1_F74G_, leading to ubiquitination and subsequent proteasomal degradation of any protein fused to mAID. Interestingly, washing of 5-Ph-IAA stops degradation and enables protein neosynthesis^8^. mAID-mCherry-iPAK4 fusion allowed us to record the precise timing of protein degradation and neosynthesis within iPAK4 fibers subjected to CATCHFiber_green_ timestamping with a match_550_[30− / 30+]_4h_ pulse sequence (**Figure 3a-d**). Timestamping-based time rescaling showed that degradation occurred during the time of exposure to 5-Ph-IAA, and that neosynthesis started after its removal (**Supplementary Text 1** and **Figure 3e**), demonstrating the suitability of CATCHFiber to record the kinetics of protein degradation and neosynthesis.

Inducing protein degradation by adding 5-Ph-IAA at different timepoints allowed us next to evaluate the precision of fiber timestamping with a match_550_[30− / 30+]_3h_ sequence (**Figure 4**). Fiber analysis showed that the degradation onset of mAID-mCherry-iPAK4 correlated with the ground truth timings of addition of 5-Ph-IAA. The average absolute mean difference to the ground truth times was 6.4 ± 2.5 min, demonstrating the high precision of CATCHFiber to report on the onset of auxin-induced degradation.

The kinetics of mAID-mCherry-iPAK4 degradation upon addition of 5-Ph-IAA were furthermore assessed in fibers using timestamping with CATCHFiber_green_ (**Figure 3f-i, and Supplementary Text 1**). The degradation of mAID fusion upon 5-Ph-IAA addition was previously measured by flow cytometry to occur with a half-time of about 62 minutes^8^. Recording the levels of mAID-mCherry-iPAK4 into fibers timestamped with a match_550_[30− / 30+]_3h_ sequence confirmed effective degradation of mAID-mCherry-iPAK4 upon addition of 5-Ph-IAA (**Figure 3f,g**). After timestamping-based time rescaling, the average time to reach 50% of degradation was determined to be 47 ± 12 min, in agreement with the reported time for the AID2 system^8^ (**Figure 3h,i**). Fitting further allowed us to refine our analysis and extract a mean decay initiation time of 9.7 ± 1.9 min and a mean half-life of 31 ± 7.3 min.

The AID2 system allows more rapid degradation than the original AID system, employing wild-type OsTIR1(WT) and large concentrations of indole-3-acetic acid (IAA)^8^. In-fiber recording of IAA-induced degradation of mAID-mCherry-iPAK4 in presence of WT OsTIR1 instead of OsTIR1_F74G_ allowed us to confirm IAA-induced degradation (**Supplementary Figure 11a-d**) and to evidence the difference in kinetics between the two systems. mCherry signal decreased to 37% of the initial levels for AID2 within 1 h of induction, while it represented still 60% in the case of AID, in agreement with a slower degradation. After 2.5 h of induction, mCherry signal decreased to 20 % for AID2, while it only reached 49% for AID (**Supplementary Figure 11e**). These results were in agreement with the half-times reported in the literature (62 minutes for AID2 and 147 minutes for AID)^8^, and demonstrate the ability of CATHFiber to distinguish such differences in kinetics.

Next, we measured the kinetics of mAID-FP-iPAK4 neosynthesis. Cells expressing mAID-mCherry-iPAK4, OsTIR1_F74G_, and CATCHFiber_green_ were pretreated with 5-Ph-IAA to induce mAID-FP-iPAK4 degradation. Then cells were subjected to a match_550_[30− / 30+]_4h_ timestamping sequence, and 5-Ph-IAA was washed away 30 minutes after timestamping start to stop degradation and enable protein neosynthesis (**Figure 3j**). A gradual increase in red fluorescence was observed along the fibers, with an onset nearing the 30-min timestamp (**Figure 3k,l**). Most fluorescence profiles reached a plateau within the 4 h timestamping period. After timestamping-based time rescaling, the protein neosynthesis half-time was determined to be 54 ± 7.5 min (**Figure 3m**), in agreement with previously reported values^8^. Incorporation of newly-synthesized mAID-FP-iPAK4 combined with CATCHFiber timestamping allows to directly access the kinetics of translation.

### Recording the cell cycle dynamics

Next, we used CATCHFiber to report on cell cycle dynamics through incorporation of geminin, a protein regulator of the cell cycle. Geminin is involved in DNA replication inhibition in the later stages of the cell cycle. Its levels increase from S to G2 phase, before its proteasomal degradation following mitosis. While human geminin localizes to the nucleus, the 1-60 truncation (hGem_1-60_) localizes both in the cell nucleus and cytosol^9^. We thus used mCherry-hGem_1-60_-iPAK4 fusion to record Geminin dynamics during the cell cycle. mCherry-hGem_1-60_-iPAK4 would accumulate in the fiber from the beginning of S-phase to the onset of mitosis, while its degradation at the end of mitosis would lead to a loss of fluorescence in the fiber. Simultaneous expression of mCherry-hGem_1-60_-iPAK4 and CATCHFiber_green_ in non-synchronized HEK293T cells submitted to a long sequence of timestamping enabled to flag different dynamics in cell-cycle evolution (**Figure 5a-c**). iPAK4 fibers showed varying levels of red fluorescence over time, with phases of complete disappearance, followed by gradual increase (**Figure 5d,f**,**h**). Using CATCHFiber, we measured the durations of hGem_1-60_ increase phases to be a few hours long (**Figure 5e,g,i)**. We identified cells that have undergone division during the course of timestamping through a sudden drop of red fluorescence in the temporal intensity profile. Noteworthily, in-fiber long-term recording enabled us to identify fibers for which the assumption of linear growth was not met (**Supplementary Figure 12**). CATCHFiber enabled us to measure the duration of expression and degradation of geminin, and identify cell cycle steps.

### Recording kinase activity

Next, we investigated whether we could record the activity of a protein kinase involved in cell signaling. We targeted cyclin-dependent kinase 2 (CDK2) – a key cell-cycle regulator whose activity can be detected using the cellular localization of its substrate DNA helicase B (DHB)^10^. DHB-FP localizes to the nucleus in the absence of CDK2 activity, while phosphorylation upon high CDK2 activity leads to translocation to the cytoplasm, enabling to follow CDK2 activity^10^.

We used a DHB-mVenus-iPAK4 fusion in combination with CATCHFiber_red_ to record CDK2 activity into fibers. Time-lapse fluorescence microscopy showed that DHB-mVenus-iPAK4 behaved similarly as the original CDK2 sensor with a half-life of nuclear translocation upon CDK2 inhibition of 6.4 ± 0.5 min (**Supplementary Figure 13**), in agreement with the 6.8 min reported for the original sensor^10^. Then, we used DHB-mVenus-iPAK4 to record the activity of CDK2 into iPAK4 fibers. Fiber timestamping was performed using CATCHFiber_red_ and a match_550_[30− / 30+]_4h_ sequence (**Figure 6**). We temporally modulated the activity of CDK2 by adding CDK1/2 inhibitor III. The inhibitor was added 30 min after timestamping sequence start, and washed out 1 h later (**Figure 6a-c**). The resulting fibers displayed a decrease in fluorescence signal in the green channel matching the 30-minute timestamp, followed by an increase, in agreement with the timings of addition and removal of the CDK1/2 inhibitor III (**Figure 6d,e**). Analysis of a large number of fibers allowed us to evaluate the dynamics of CDK2 activity upon addition of CDK1/2 inhibitor III. We extracted an average half-life decay of 24 ± 5.4 min, and an average decay initiation time of 12 ± 3.1 minutes (**Figure 6f**). These values are slightly higher than the values extracted from analyzing the time-lapses. This difference can stem from the difference in the measured information. The analysis of the time-lapses uses the cytosol-to-nucleus fluorescence ratio as a proxy of CDK2 activity, while the fluorescence along the fiber only accounts for the variations in cytosolic DHB-mVenus-iPAK4. Nevertheless, this set of experiments convincingly shows that CATCHFiber enables to estimate the temporal dynamics of CDK2 activity.

**Figure 6.**
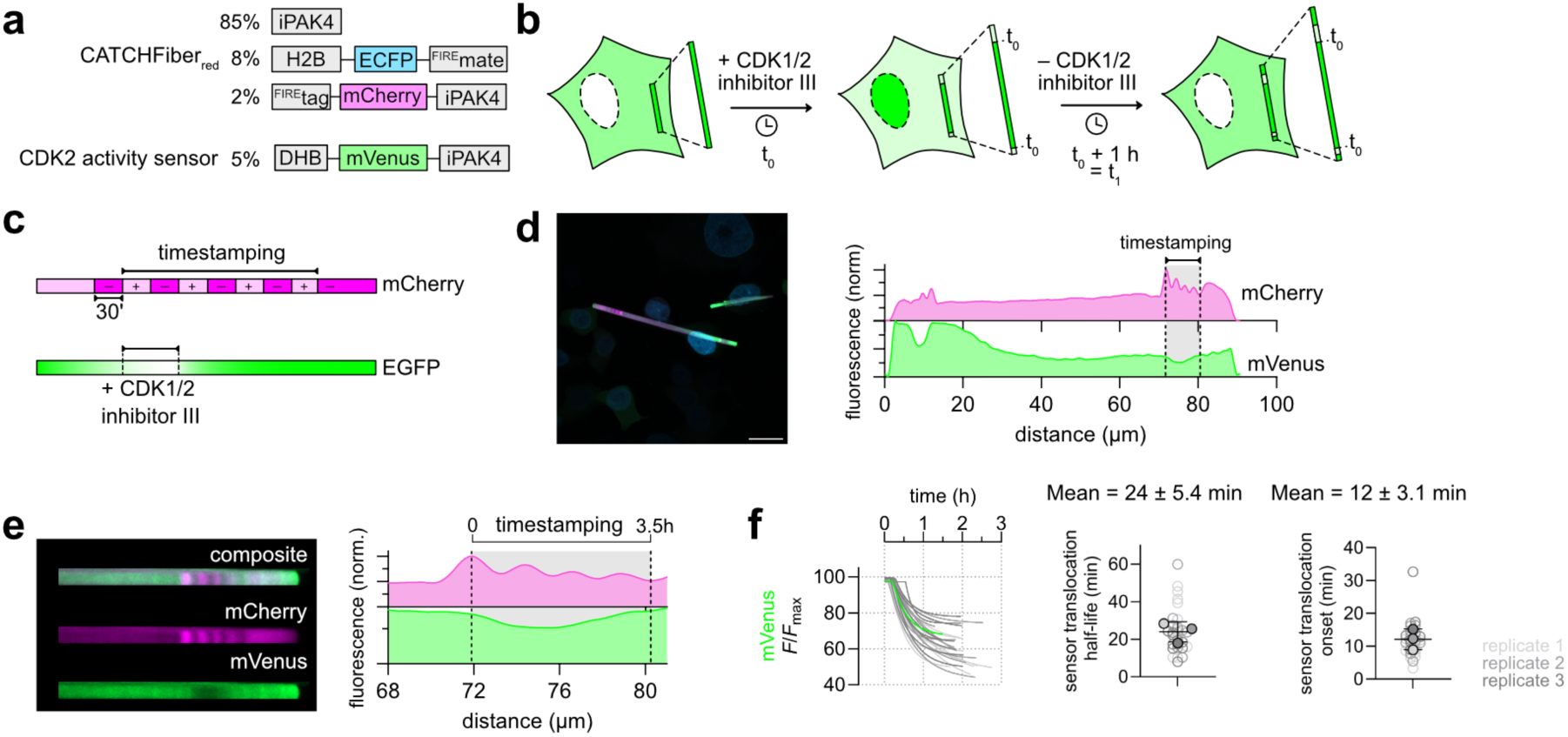
CATCHFiber enables the recording of CDK2 activity. **a** Plasmids used. **b-f** HEK293T cells expressing CATCHFiber_red_ were subjected to the timestamping sequence match_500_[30− / 30+]_4h_ and CDK1/2 inhibitor III (10 µM) was added along with the first match_550_ addition and washed 1h later. **d** A representative micrograph as well as the fluorescence intensity plot along the displayed fiber is shown. Scale bar 20 µm. The timestamping period is shown as a grey highlight in the full fluorescence intensity plot. A straightened image of the full fiber is also shown. The dashed black lines positions were normalized to 0 and 3.5. **e** Zoomed-in image of the fiber and the corresponding intensity plot (smoothed). The results are representative of n = 29 fibers from three replicates. **f** Fitted temporal evolution of mVenus intensity from all 29 fibers imaged in this experiment after temporal rescaling and fitting using a one phase-decay model with an initial plateau. Each curve is color-coded according to the replicate the fiber comes from. In green is the fitted temporal evolution of the fiber shown in (**d-e**). For each fitted curve, are extracted and plotted the degradation half-life and onset. In each plot, each fiber is color-coded according to the biological replicate it came from. The solid circles correspond to the mean of each biological replicate. The black line represents the mean ± SD of the three replicates (n = 29 fibers).

## DISCUSSION

Protein-based fiber systems enable to record intracellular events in space, opening interesting perspectives to store cell history. Fiber snapshots can now tell similar stories to long-term multi-field of view time-lapses. Large-scale single-cell recording of transcriptional activity was previously achieved with the iPAK4 ticker tapes^3,6^, as well as the XRI^2^ and CytoTape^5^ systems. Here, we present CATCHFiber, a chemogenetic timestamping strategy for precise tracing of cell history. CATCHFiber enables to place fiducial timestamps on iPAK4 fibers, as narrowly-spaced as 30 minutes, overcoming the limited temporal resolution of previously reported HaloTag-based timestamps^3^. CATCHFiber relies on chemically-induced dimerization (CID) to reversibly control the cellular localization of fluorescent timestamping monomers, yielding user-controlled alternating dark and bright marks along the fibers. Timestamping using a single fluorescent protein rather than multiple dyes advantageously frees detection channels for multiplexed recordings.

The duration of CATCHFiber-induced timestamps can be tuned to match the temporality of the recorded event. Timestamps can be spaced by as short as 30 minutes, up to 12 hours for the longest experiments reported here, enabling to cover a wide range of biological intracellular timescales. Closely-spaced timestamps enable to correct for fluctuations in fiber linear growth rate, especially during long temporal windows. Given the kinetics of CATCHFIRE used for timestamping control, one can imagine even more closely spaced timestamps, although the growth rate of iPAK4 fibers, reported to be about 1.5 µm/h^3^, will probably limit timestamps spacing to 15 minutes at best, as such timestamps would be spaced by 0.4 µm, which is at the resolution limit of fluorescence microscopy.

So far, iPAK4 ticker-tapes, XRI and CytoTape systems have only been used to record gene expression. Here we show that the increased temporal resolution of CATCHFiber expands the possibilities of fiber-based recording to faster cellular processes. Timestamping strategy based on HaloTag enabled to place time stamps every 2 hours at best, preventing the precise recording of fast events occurring at shorter timescales. Here, we successfully assessed the precise kinetics of events with sub-hour half-life such as inducible protein degradation with the AID2 system, and could evidence the kinetics difference observed with the original AID system^8^. Leveraging the reversibility of the AID2 degradation system, we also evaluated the rapid kinetics of neosynthesis (i.e. translation) following auxin-derivative washout. The approach could be used in the future to evaluate with single-cell resolution the performance of chemical inducers of degradation (e.g. molecular glues, PROTACs) for biological and therapeutical applications.

The high temporal resolution of CATCHFiber timestamping also allowed us to precisely record the activity of protein kinases such as CDK2 using a translocation-based kinase sensor. In-fiber recording enables to uncover kinase activity on a large scale conserving single-cell precision, opening interesting perspectives for screening potent kinase inhibitors and study their dynamics. This strategy is general and could be expanded in the future to the design of recorders for other enzymes and cellular processes using existing translocation-based sensors.

Finally, we also showed that CATCHFiber could be used to access the dynamics of intracellular events such as cell division, using the degradation and synthesis of hGeminin1-60 for delineating cell cycle stages. The use of iPAK4 fibers for recording such events is however not optimal as their rigidity impedes complete cytokinesis and thus influences cellular division. We anticipate that CATCHFiber timestamping could be combined with more flexible fiber scaffolds as XRI and CytoTape^5^ or novel assemblies like GEMINI, which enabled to overcome the limitations linked to the use of linear protein assemblies for recording^11^.

Future efforts will involve the automation of the match molecule addition and washouts – currently manually handled – to facilitate the deployment of CATCHFiber in different biological contexts. Automation will improve reproducibility and scalability through parallelized timestamping.

In conclusion, CATCHFiber enables in-fiber timestamping with unprecedented temporal resolution enabling to record biological events from sub-30 min to several hours timescales, enabling to expand the application of fiber-based recording beyond transcriptional activity. We envision that the high temporal precision it offers will make it an interesting system for tracing the history of cells and uncovering the dynamics of various cellular processes.

## MATERIALS & METHODS

### General

Commercially available reagents were used as obtained. The synthesis of match_550_ (a.k.a HBR-2,5DM) was previously reported^12^. This molecule is commercially available from the Twinkle Factory under the name match_550_.

### Biology

The presented research complies with all relevant ethical regulations.

### General

Synthetic oligonucleotides used for cloning were purchased from Integrated DNA Technology. PCR reactions were performed with Q5 polymerase (New England Biolabs) in the buffer provided. PCR products were purified using QIAquick PCR purification kit (QIAGEN). DNase I, T4 ligase, fusion polymerase, Taq ligase and Taq exonuclease were purchased from New England Biolabs and used with accompanying buffers and according to the manufacturer’s protocols. Isothermal assemblies (Gibson Assembly) were performed using a homemade mix prepared according to previously described protocols^13^. Small-scale isolation of plasmid DNA was conducted using a QIAprep miniprep kit (QIAGEN) from 2-3 mL overnight bacterial culture supplemented with appropriate antibiotics. Large-scale isolation of plasmid DNA was conducted using the QIAprep maxiprep kit (QIAGEN) from 150 mL overnight bacterial culture supplemented with appropriate antibiotics. All plasmid sequences were confirmed by Sanger sequencing with appropriate sequencing primers. All the plasmids used in this study are listed in **Supplementary Tables 1 and 2**, as well as their DNA sequences and ratio of utilization.

### Cloning

DL015:CMV::iPAK4 was a gift from Adam Cohen (Addgene plasmid # 177880 ; http://n2t.net/addgene:177880 ; RRID:Addgene_177880)^3^. DL016:CMV::eGFP-iPAK4 was a gift from Adam Cohen (Addgene plasmid # 177881 ; http://n2t.net/addgene:177881 ; RRID:Addgene_177881)^3^. DL146_plenti_TRE_GFP-iPAK4 was a gift from Adam Cohen (Addgene plasmid # 187446; http://n2t.net/addgene:187446 ; RRID:Addgene_187446)^3^. DHB-mVenus was a gift from Tobias Meyer & Sabrina Spencer (Addgene plasmid # 136461 ; http://n2t.net/addgene:136461; RRID:Addgene_136461)^10^.

The plasmids used in this study have been generated using isothermal Gibson assembly.

Plasmid pAG1903 for mammalian expression of FLAG-iPAK4 under the control of a CMV promoter was obtained by replacing the sequence coding for pFAST by FLAG-iPAK4 amplified from addgene plasmid #177880 in the plasmid pAG654 enabling mammalian expression of pFAST.

Plasmid pAG1904 for mammalian expression of EGFP-SGGS-iPAK4 was obtained by replacing the sequence coding for pFAST by EGFP-SGGS-iPAK4 amplified from addgene plasmid #177881 in the plasmid pAG654 enabling mammalian expression of pFAST.

Plasmid pAG1907 (respectively pAG1908) for mammalian expression of mCherry-SGGS-iPAK4 (respectively emiRFP670-SGGS-iPAK4) was obtained by replacing the sequence of EGFP by the sequence of mCherry (respectively emiRFP670) amplified from plasmid pAG1160 coding for mCherry-^FIRE^tag (respectively amplified from plasmid pAG1335 coding for emiRFP670-P2A-EGFP) in the plasmid pAG1904.

Plasmid pAG1910 for mammalian expression of ^FIRE^tag- SGGGGSGG-mCherry-iPAK4 was obtained by replacing the sequence of mCherry by ^FIRE^tag-SGGGGSGG-mCherry amplified from plasmid pAG1209 encoding ^FIRE^tag-SGGGGSGG-mCherry in the plasmid pAG1907.

Plasmid pAG1909 (respectively pAG1911) for mammalian expression of ^FIRE^tag-SGGGGSGG-EGFP-iPAK4 (respectively ^FIRE^tag-SGGGGSGG-emiRFP670-iPAK4) was obtained by replacing mCherry by EGFP (respectively emiRFP670) amplified from the plasmid pAG1904 encoding EGFP-SGGS-iPKA4 (respectively from the plasmid pAG1908 encoding emiRFP670-SGGS-iPAK4) in the plasmid pAG1910.

Plasmid pAG2054 (respectively pAG2055) for mammalian expression of mAID-AS-EGFP-SGGS-iPAK4 (respectively mAID-AS-mCherry-SGGS-iPAK4) was obtained by introducing the sequence of mAID amplified from the plasmid pAG001 encoding Tir1-9cMyc-IRES-MCS-HA-EGFP-AID-NLS in the plasmid pAG1904 (respectively pAG1907).

Plasmid pAG2062 for mammalian expression of OsTIR1-cMyc was obtained by replacing the sequence of H2B-pFAST in the plasmid pAG657 coding for H2B-pFAST-cMyc by the sequence of OsTIR1 amplified from plasmid pAG001 encoding Tir1-9cMyc-IRES-MCS-HA-EGFP-AID-NLS.

Plasmid pAG2063 for mammalian expression of OsTIR1_F74G_-cMyc was obtained by introducing the mutation F74G in the sequence of OsTIR1 in the plasmid pAG2062 encoding OsTIR1-cMyc.

Plasmid pAG2064 for mammalian expression of rtTA3-cMyc was obtained by replacing the sequence coding for pFAST by the sequence coding for rtTA3 in the plasmid pAG654 enabling mammalian expression of pFAST.

Plasmid pAG2099 for mammalian expression of DHB-mVenus-iPAK4 was obtained by introducing the sequence DHB-mVenus amplified from addgene plasmid #136461 coding for DHB-mVenus in the plasmid pAG1903 encoding FLAG-iPAK4.

Plasmid pAG2101 for mammalian expression of mCherry-hGem_(1-60)_-iPAK4 was obtained by introducing the sequence of hGem_(1-60)_ amplified from the plasmid pAG477 encoding cMyc-redFAST-Cdt1_(30-120)_-P2A-greenFAST-hGem_(1-60)_ in the plasmid pAG1907 coding for mCherry-SGGS-iPAK4.

### Cell culture

HEK293T (ATCC CRL-3216) cells were cultured in Dulbecco’s modified Eagle medium (DMEM) supplemented with phenol red,10% (vol/vol) FBS and 1% (vol/vol) penicillin–streptomycin at 37 °C in a 5% CO_2_ atmosphere. For imaging, cells were seeded in μDish ibidi (Biovalley) coated with poly-L-lysine. Cells were transiently transfected using Genejuice (Merck) according to the manufacturer’s protocols 1-2 days prior to imaging.

*Timestamping procedure* For CATCHFiber implementation in mammalian cells, HEK293T cells were transfected as described above with the corresponding plasmids at the indicated ratios for each experiment (**Supplementary Table 1**) in the presence of 10 µM of match_550_ directly added in the culture medium (DMEM + 10% (vol/vol) FBS + 1% (vol/vol) penicillin–streptomycin). The following day, the cells were subjected to a sequence of match_550_ washout and addition as described for each experiment. Match_550_ washout was performed through removal of medium, and subsequent 5 washes with a total of 6 mL of DPBS. Addition of match_550_ was performed through replacing match-free DMEM with DMEM supplemented with 10 µM of match_550_. Cells were returned to the incubator during the long incubations in the presence or in the absence of match_550_.

*Induction of expression* For induction of expression using the tetON system, the culture medium was supplemented with 10% (vol/vol) Tet-System approved FBS (A4736401, Gibco) and 1% (vol/vol) penicillin–streptomycin. Induction of EGFP expression was performed through addition of doxycycline (doxycycline hyclate, D5207, Sigma-Aldrich) directly into the culture medium (final concentration 2 µg/mL) at the desired timing.

*Induction of degradation* Targeted protein degradation using the AID system was performed through addition of 3-Indoleacetic acid (I3750, Sigma-Aldrich) directly into the culture medium (final concentration 100 µM) at the desired timing. Targeted protein degradation using the AID2 system was performed through addition of 5-Phenyl-indole-3-acetic acid (SML3574, Sigma Aldrich) directly into the culture medium (final concentration 1 µM) at the desired timing.

*Recording of kinase activity* CDK2 activity was inhibited through addition of CDK1/2 Inhibitor III (217714, Sigma-Aldrich) directly into the culture medium (final concentration 10 µM) at the desired timing.

### Live cell imaging

The confocal micrographs of mammalian cells were acquired on an inverted Zeiss LSM 980 Laser Scanning Microscope equipped with a plan apochromat 63× /1.4 NA oil immersion objective and a plan-apochromat 20× /0.8 dry. The wavelengths used for excitation were 445 nm for ECFP, 488 nm for EGFP, mVenus and match_550_, 561 for mCherry and 639 for emiRFP670. ZEN software was used to collect the data. Icy and Fiji softwares were used to analyze the data.

### Image and data analysis

The different fields of view displayed in this work were acquired either as multicolor snaps in the case of integrally in-focus fibers or as multicolor z-stacks with a range covering the full fiber length. In the case of z-stacks, the maximum intensity projection was used to retrieve the fluorescence profiles along the full fiber length. Fluorescence intensity profiles were obtained through manually-defined line regions of interest with widths matching the fiber width using the plot profile tool, and saved for each channel of interest for the fibers. The straighten tool on the selected fiber line ROI was used exclusively for display purposes to align the fibers horizontally.

Graphpad 11 software was used to handle the extracted fluorescence intensities and plot the different graphs. For each channel, the fluorescence intensity was smoothed (4 or 9 neighbors depending on the graphs) and the first and second derivatives were computed (with smoothing at 4 neighbors). The timestamps were identified as cancellation points of the first derivative of the fluorescence intensity in the timestamping channel, and the recorded events onsets were identified as local extrema of the second derivative of the fluorescence intensity in the recording channel (see **Supplementary Text 1** and **Annex Figures 1 and 2)** for more detailed explanation).

## STATISTICS AND REPRODUCIBILITY

No sample size calculations were performed. When relevant, the sample size (n) is provided in the corresponding figure captions. Sample sizes were chosen to support meaningful conclusions. No data were excluded. The number of replicates for each individual experiments is indicated in the figure legends. All attempts at replication were successful. The experiments were not randomized. The Investigators were not blinded to allocation during experiments and outcome assessment.

## DATA AVAILABILITY

Data supporting the findings of this study are available within the article and supplementary information, and are available from the corresponding authors upon request. The plasmids developed in this study may be requested from the corresponding author.

## ACKNOWLEDGMENTS

We thank France Lam, Chloé Chaumeton and Romain Naillon from the Institut de Biologie Paris Seine imaging platform for their assistance with imaging. We particularly acknowledge Jean-François Gilles for his help with the data extraction macro on Fiji. This work has been supported by the Agence Nationale de la Recherche (ANR-23-CE44-0014-01 CATCHFIRE), the Institut Universitaire de France, and the Dynamic Imaging program of the Chan-Zuckerberg Initiative DAF (grant number 2023-321185), an advised fund of Silicon Valley Community.

## AUTHOR CONTRIBUTIONS

L.E.H and A.G. designed the overall project and wrote the paper. L.E.H and A.G. designed the experiments. L.E.H. performed the experiments. L.E.H and A.G analyzed the experiments.

