## Supporting Information for "Chemogenetic timestamping for the precise tracing of cell history into protein assemblies"

### Content

**Supplementary Text 1**

**Supplementary Figures 1-13**

**Supplementary Tables 1-2**

**Supplementary text 1 – Description of the analysis process**

CATCHFiber consists of a large excess of iPAK4 monomers which drives fiber formation, combined with $^{FIRE}$tag-FP-iPAK4 – acting as a timestamping monomer –, H2B-ECFP-$^{FIRE}$mate – the nucleus anchoring moiety – and the small fluorogenic ligand $match_{550}$. In HEK293T cells expressing CATCHFiber, $^{FIRE}$tag-FP-iPAK4 traffics to the cell nucleus and associates with H2B-ECFP-$^{FIRE}$mate upon addition of $match_{550}$, which reduces its cytosolic concentration. Consequently, the fiber grows in the absence of this fluorescent monomer during incubation with $match_{550}$, leaving a dark mark. Conversely, $match_{550}$ washout leads to an increase in the cytosolic concentration of $^{FIRE}$tag-FP-iPAK4, which allows its incorporation in the iPAK4 fibers, leading to a fluorescent mark. Consequently, match addition and washout results in alternating bright and dark bands on the fiber. In parallel, an event of interest triggered or occurring during the course of timestamping leaves an additional fluorescent mark.

The CATCHFiber-induced timestamps on iPAK4 fibers allow to spatially-encode time. In order to leverage these timestamps to uncover the kinetics of the recorded events, we developed a general analysis workflow, illustrated in **Annex Figures 1 and 2**. Briefly, the fluorescence intensity along the fiber in the channels of interest is retrieved, smoothed, and the first derivative of the curve is computed. In the timestamping channel, the fluorescence intensity plot shows alternating maxima and minima, correlating with $match_{550}$ washouts and additions. These extrema can be precisely determined by finding the cancellation points of the first derivative of the fluorescence intensity curve. For each experiment, the conversion of the spatial information to temporal information was either performed using the timing of the first match addition and the last washout, or by using the information from all timestamps (meaning all cancellation points of the $1^{st}$ derivative).

---

**Figure 2 – Analysis of the timestamping on both iPAK4 fiber ends**

*Sequence* – $match_{550}[30–/\ 30+]_{4h}$

*Position normalization* – the position was normalized between the timestamps t = 0.5 h (reset to t = 0 h) and timestamps t = 4 h (reset to t = 3.5 h). The positions of the timestamps were determined using the cancellation points of the first derivative of the fluorescence intensity profile. Note that choosing the timestamps t = 0.5 h rather than t = 0 h for position normalization simplifies the systematic detection of this point during the analysis. The maximum at t = 0.5 h is more precisely identified with the cancellation of the derivative of the intensity profile.

*First derivative* – The first derivative was calculated on smoothed fluorescence intensity (4 neighbors on each side for the curve and derivative).

*Observations* – Note that several fibers were excluded from the analysis as the pattern was unmistakably observed on the faster-growing side, but not on the slower-growing one. One possible explanation for such behavior is that, given the reported average growth rate of iPAK4 fibers in HEK293T cells (1.46 ± 0.64 μm/h[1]), in this case they have not grown enough during the 30 min period between two timestamps, leading to patterns that cannot be resolved with conventional confocal microscopy. In order to maximize the number of analyzable fibers for further event recording experiments and to benefit from the maximal pattern precision, we decided to systematically analyze the fluorescence patterns on the faster growing side.

---

**Figure 3c-e – AID2 reversibility evaluation**

*Sequence* – $\text{match}_{550}[30-/\ 30+]_{4h}$

*Position normalization* – The position was normalized between the timestamps t = 0.5 h (reset to t = 0 h) and t = 4h (reset to t = 3.5 h). The positions of the timestamps were determined using the cancellation points of the first derivative of the fluorescence intensity profile.

*First derivative* – The first derivative was calculated on smoothed fluorescence intensity (9 neighbors on each side for the curve and 4 for the derivative).

*Temporal rescaling* – The information of all the timestamps was used for precise rescaling of the reporter signal. For temporal rescaling, the distance between two consecutive 30-min timestamps was normalized to 0.5, allowing to directly convert the position into time.

Similar analysis applies for **Supplementary Figure 10e**.

---

**Figure 3f-l – Degradation kinetics evaluation**

*Sequence* – $\text{match}_{550}[30-/\ 30+]_{3h}$

*Position normalization* – The position was normalized between timestamps t = 0.5 h (reset to t = 0 h) and t = 3 h (reset to t = 2.5 h). The positions of the timestamps were determined using the cancellation points of the first derivative of the fluorescence intensity profile.

*First derivative* – The first derivative was calculated on smoothed fluorescence intensity (9 neighbors on each side for the curve and 4 for the derivative).

*Temporal rescaling* – The information of all the timestamps was used for precise rescaling of the reporter signal. For temporal rescaling, the distance between two consecutive 30-min timestamps was normalized to 0.5, allowing to directly convert the position into time.

*Degradation half-time* – After temporal rescaling, mCherry signal was normalized between 0 and maximal fluorescence $F_{max}$. For each fiber from each replicate, the time positions to reach $F = F_{max}/2$ is reported.

*Data fit* – The degradation curves were fitted with a one phase decay model with an initial plateau. (Y= IF( X<X0, Y0, Plateau + (Y0 – Plateau)*exp(–K*(X–X0)))), and the values of X0 (onset) and ln(2)/K (half-life) are reported.

Similar analysis applies for **Supplementary Figure 10j**.

---

**Figure 3j-m – Neosynthesis kinetics evaluation**

*Sequence* – $\text{match}_{550}[30-/\ 30+]_{4h}$

*Position normalization* – The position was normalized between timestamps t = 0.5 h (reset to t = 0) and t = 4 h (reset to t = 3.5h). The positions of the timestamps were determined using the cancellation points of the first derivative of the fluorescence intensity profile.

*First derivative* – The first derivative was calculated on smoothed fluorescence intensity (9 neighbors on each side for the curve and 4 for the derivative).

*Temporal rescaling* – The information of all the timestamps was used for precise rescaling of the reporter signal. For temporal rescaling, the distance between two consecutive 30-min timestamps was normalized to 0.5, allowing to directly convert the position into time.

*Neosynthesis half-time* – After temporal rescaling, mCherry signal is normalized between minimal fluorescence $F_{min}$ and maximal fluorescence $F_{max}$. For each fiber from each replicate the time positions to reach $(F - F_{min}) / (F_{max} - F_{min}) = 50$ % (half-time of degradation) is reported.

Similar analysis applies for **Supplementary Figure 10n**.

---

**Figure 4 – Degradation onset evaluation**

*Sequence* – $match_{550}[30-/30+]_{3h}$

*Position normalization* – The position was normalized between timestamps t = 0.5 h (reset to t = 0) and t = 3 h (reset to t = 2.5 h). The positions of the timestamps were determined using the cancellation points of the first derivative of the fluorescence intensity profile.

*First derivative* – The first derivative was calculated on smoothed fluorescence intensity (9 neighbors on each side for the curve and 4 for the derivative).

*Temporal rescaling* – The information of all the timestamps was used for precise rescaling of the reporter signal. For temporal rescaling, the distance between two consecutive 30-min timestamps was normalized to 0.5, allowing to directly convert the position into time.

*Degradation onset* – After temporal rescaling, mCherry signal was normalized between 0 and maximal fluorescence $F_{max}$. The degradation onset was identified through identification of local minima in the fluorescence intensity second derivative curve (smoothing equal to 4). For the condition where 5-Ph-IAA was added at t = 0 h, as some decays initiated without measurable onset, the corresponding curves were omitted from this calculation.

---

**Figure 5 – Recording of the cell cycle dynamics using CATCHFiber**

*Sequence* – $match_{550}[30-/120+]_{5h}$ then $match_{550}[30-/730+]_{12h40}$

*Position normalization* – The position was normalized between t = 0.5 h (reset to t = 0) and t = 17.7 h (reset to t = 17.2 h). The positions of the timestamps were determined using the cancellation points of the first derivative of the fluorescence intensity profile.

*First derivative* – The first derivative was calculated on smoothed fluorescence intensity (9 neighbors on each side for the curve and 4 for the derivative).

*Temporal rescaling* – The information of all the timestamps was used for precise rescaling of the reporter signal. For temporal rescaling, the distance between two consecutive timestamps was normalized to reflect the time in between the two timestamps, allowing to directly convert the position into time.

Similar analysis applies for **Supplementary Figure 12**.

---

**Figure 6 – Recording of kinase activity in mammalian cells using CATCHFiber**

*Sequence* – $match_{550}[30- / 30+]_{4h}$

*Position normalization* – The position was normalized between t = 0.5 h (reset to t = 0) and t = 4 h (reset to t = 3.5h). The positions of the timestamps were determined using the cancellation points of the first derivative of the fluorescence intensity profile.

*First derivative* – The first derivative was calculated on smoothed fluorescence intensity (9 neighbors on each side for the curve and 4 for the derivative).

*Temporal rescaling* – The information of all the timestamps was used for precise rescaling of the reporter signal. For temporal rescaling, the distance between two consecutive 30-min timestamps was normalized to 0.5, allowing to directly convert the position into time.

*Data fit* – After temporal rescaling, the degradation curves were fitted with a one phase decay model with an initial plateau. (Y= IF( X<X0, Y0, Plateau + (Y0 – Plateau)*exp(–K*(X–X0)))), and the values of X0 (onset) and ln(2)/K half-time are reported.

*Observation* – mVenus protein seems to accumulate more on one end of the fibers, as evidenced by higher levels of green fluorescence on one side compared to the other. Using the timestamps information, it seems that almost systematically, this end is the slower-growing end. We used nevertheless the faster-growing side because of easier analysis of the timestamps, although the variations of the green fluorescence were less pronounced.

---

**Supplementary Figure 7 – Analysis of the pulse sequence duration on the timestamps**

*Sequence* – $match_{550}[30- / 30+]_{4h,}$ $match_{550}[60- / 60+]_{4h}$ and $match_{550}[120- / 120+]_{4h}$

*Position normalization* – the position was normalized between the timestamps t = 0 h and timestamps t = 4 h. The positions of the timestamps were determined using the cancellation points of the first derivative of the fluorescence intensity profile.

*First derivative* – The first derivative was calculated on smoothed fluorescence intensity (4 neighbors on each side for the curve and derivative).

---

**Supplementary Figure 9 – Recording gene activation with CATCHFiber**

*Sequence* – $match_{550}[30- / 60+]_{4.5h}$. In this pulse sequence, the duration of the – match incubation is shorter than the + match incubation. This ensures that the transitions are sharp enough and easier to delineate unmistakably, in comparison with an equivalent $match_{550}[60- / 60+]_{4.5h}$.

*Position normalization* – The position was normalized between the timestamps t = 0.5 h (reset to t = 0 h) and t = 4.5 h (reset to t = 4 h). The positions of the timestamps were determined using the cancellation points of the first derivative of the fluorescence intensity profile.

*First derivative* – The first derivative was calculated on smoothed fluorescence intensity (9 neighbors on each side for the curve and 4 for the derivative).

*Temporal rescaling* – The information of all the timestamps was used for precise rescaling of the reporter signal. For temporal rescaling, the distance between two consecutive timestamps was normalized to 0.5 or 1 depending if the timestamps are spaced by 30 min or 60 min, respectively, allowing to directly convert the position into time.

---

**Supplementary Figure 11 – Degradation with AID kinetics evaluation**

*Sequence* – $match_{550}[30-/ 30+]_{3h}$

*Position normalization* – The position was normalized between timestamps t = 0.5 h (reset to t = 0 h) and t = 3 h (reset to t = 2.5 h). The positions of the timestamps were determined using the cancellation points of the first derivative of the fluorescence intensity profile.

*First derivative* – The first derivative was calculated on smoothed fluorescence intensity (9 neighbors on each side for the curve and 4 for the derivative).

*Temporal rescaling* – The information of all the timestamps was used for precise rescaling of the reporter signal. For temporal rescaling, the distance between two consecutive 30-min timestamps was normalized to 0.5, allowing to directly convert the position into time.

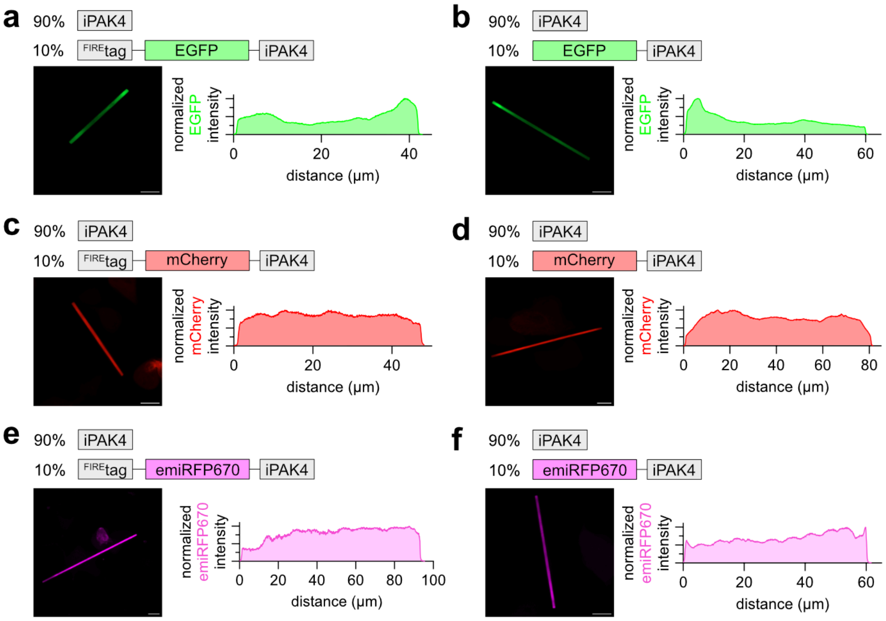


**Supplementary Figure 1. Incorporation of $^{FIRE}$tag-FP-iPAK4 and FP-iPAK4 into iPAK4 fibers.** HEK293T cells were transfected with plasmids encoding iPAK4 (90%) together with plasmids (10%) encoding $^{FIRE}$tag-EGFP-iPAK4 (**a**), EGFP-iPAK4 (**b**), $^{FIRE}$tag-mCherry-iPAK4 (**c**), mCherry-iPAK4 (**d**), $^{FIRE}$tag-emiRFP670-iPAK4 (**e**) or emiRFP670-iPAK4 (**f**), and imaged the day after transfection. Representative micrographs of fluorescently labeled fibers and the corresponding fluorescence intensity plots (smoothed) are shown. **a,c,e** Representative results from three replicates: (**a**) n = 36 fibers, (**c**) n = 25 fibers, (**e**) n = 32 fibers (See also **Supplementary Figure 2**). **b,d,f** Representative results from two replicates: (**b**) n = 15 fibers, (**d**) n = 14 fibers, (**f**) n = 18 fibers (See also **Supplementary Figure 3**). Scale bar 10 µm

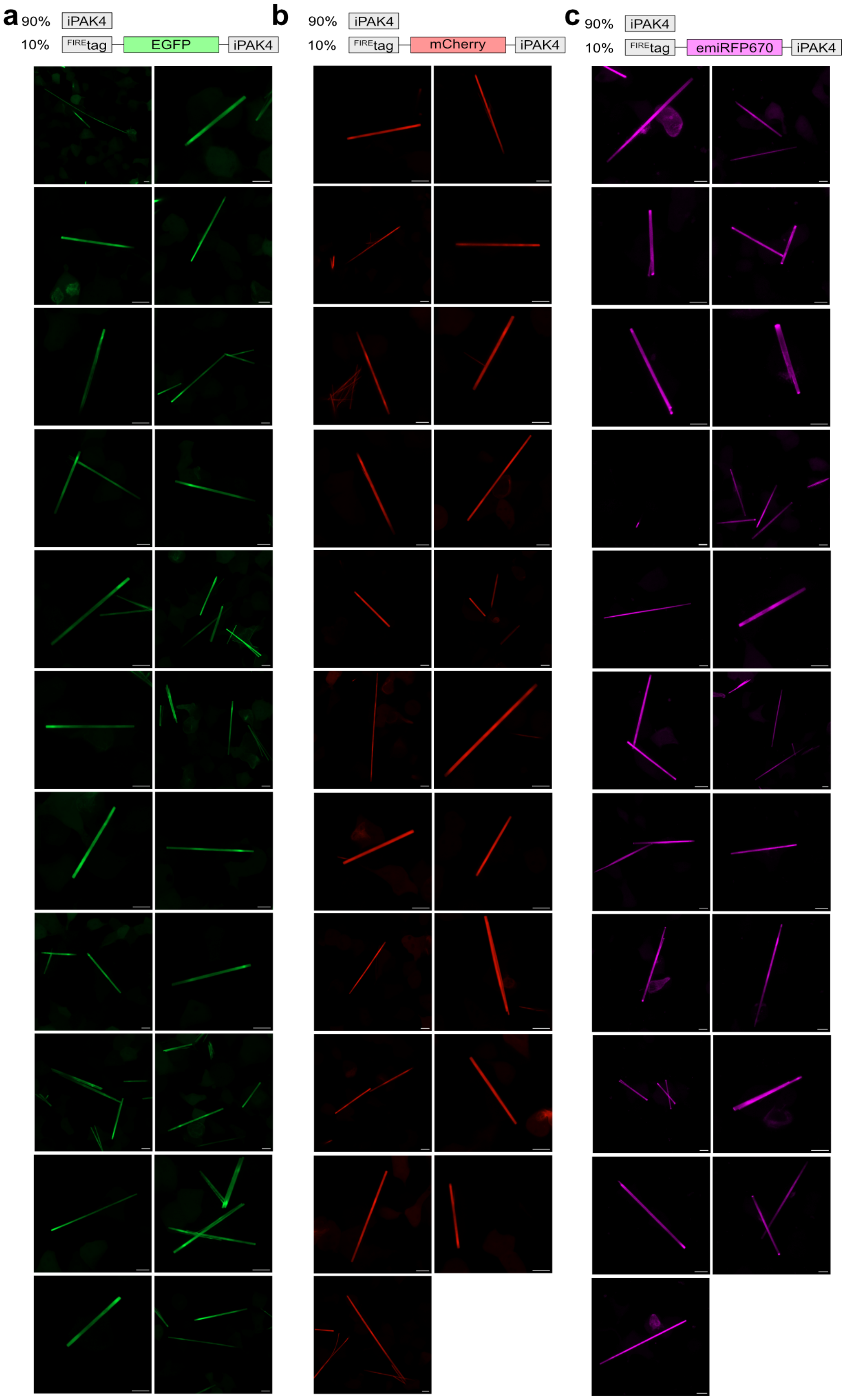


**Supplementary Figure 2. Incorporation of FIREtag-FP-iPAK4 into iPAK4 fibers.** HEK293T cells were transfected with plasmids encoding iPAK4 (90%) together with plasmids (10%) encoding FIREtag-EGFP-iPAK4 (**a**), FIREtag-mCherry-iPAK4 (**b**), FIREtag-emiRFP670-iPAK4 (**c**) and imaged the day after transfection. Are shown micrographs of fluorescently labeled fibers from three replicates: (**a**) n = 36 fibers, (**c**) n = 25 fibers, (**d**) n = 32 fibers. Scale bar 10 μm

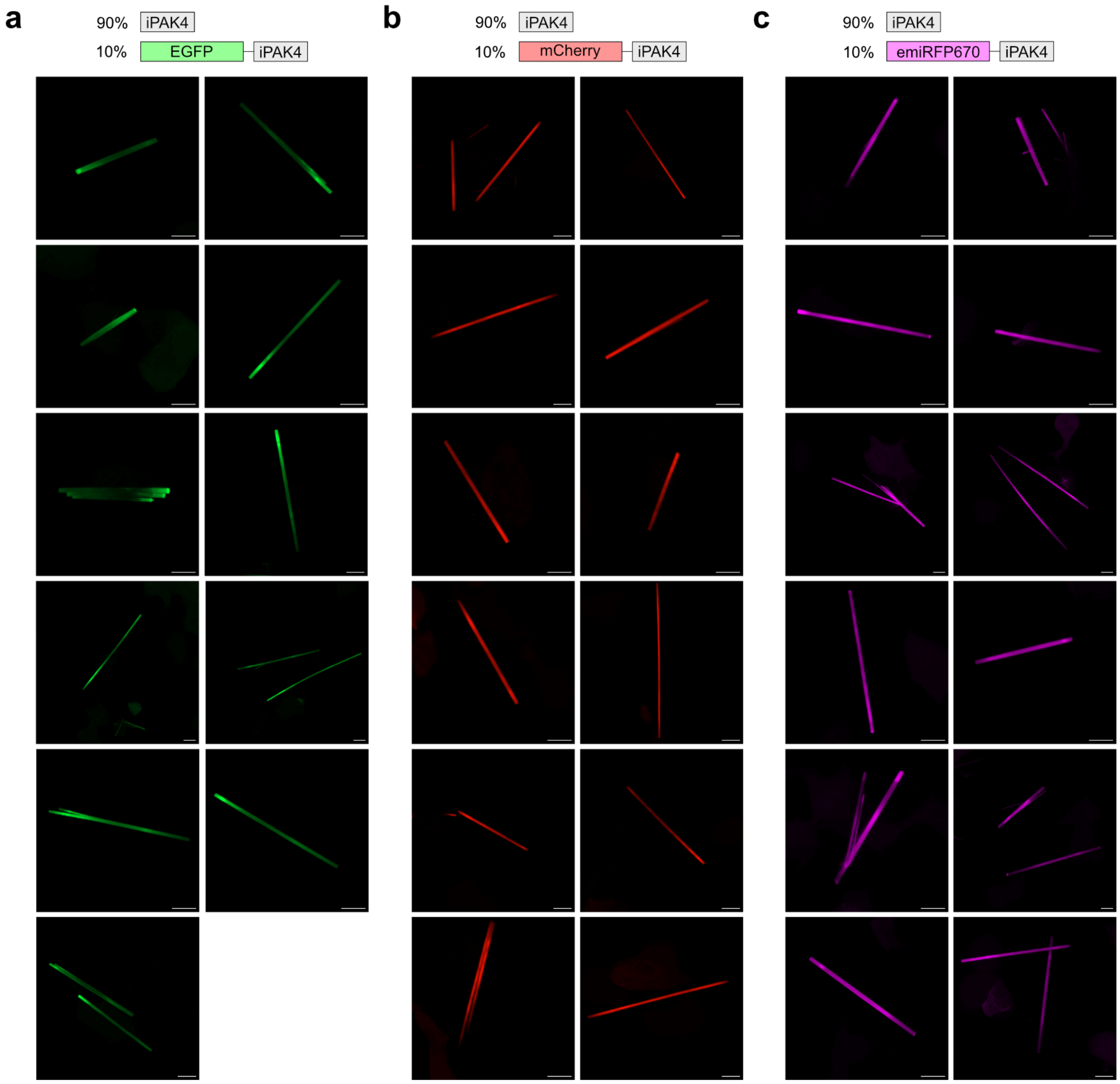


**Supplementary Figure 3. Incorporation of FP-iPAK4 into iPAK4 fibers.** HEK293T cells were transfected with plasmids encoding iPAK4 (90%) together with plasmids (10%) encoding EGFP-iPAK4 (**a**), mCherry-iPAK4 (**b**), emiRFP670-iPAK4 (**c**), and imaged the day after transfection. Are shown micrographs of fluorescently labeled fibers from two replicates: (**a**) n = 15 fibers, (**b**) n = 14 fibers, (**c**) n = 18 fibers. Scale bar 10 µm

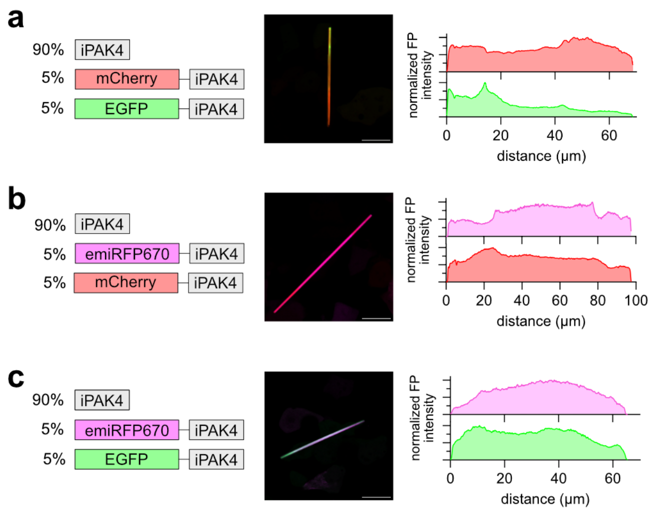


**Supplementary Figure 4. Incorporation of two fluorescent proteins into iPAK4 fibers.** HEK293T cells were transfected with plasmids encoding iPAK4 (90%) together with plasmids (5%) encoding (**a**) mCherry and EGFP, (**b**) emiRFP670 and mCherry, or (**c**) emiRFP670 and EGFP. Representative micrographs of fluorescently labeled fibers from two replicates (n > 50 fibers) and the corresponding fluorescence intensity plots (smoothed) are shown (see also **Supplementary Figure 5**). Scale bar 20 µm.

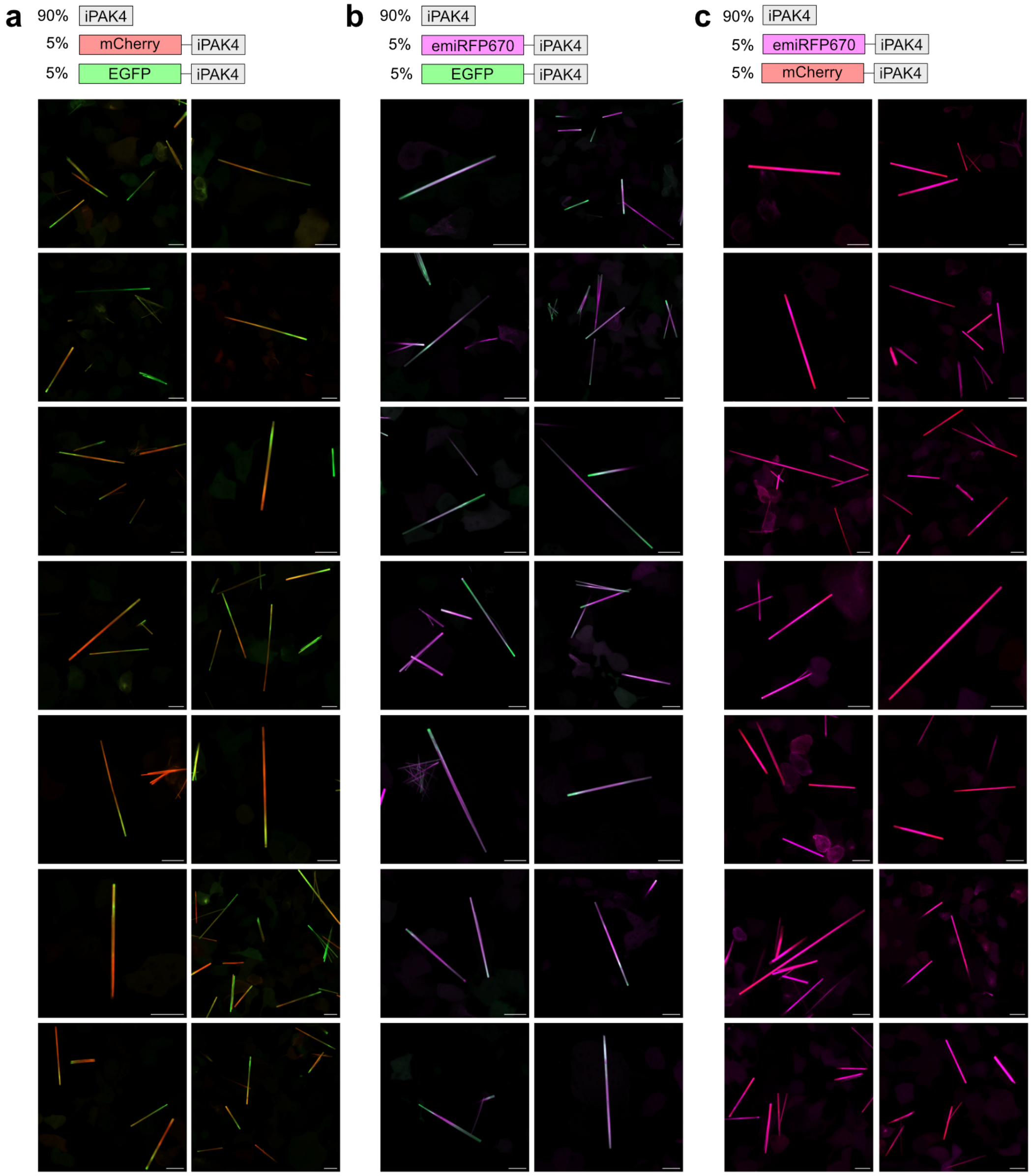


**Supplementary Figure 5. Incorporation of two fluorescent proteins into iPAK4 fibers.** HEK293T cells were transfected with plasmids encoding iPAK4 (90%) together with plasmids (5%) encoding (**a**) mCherry and EGFP, (**b**) emiRFP670 and mCherry, or (**c**) emiRFP670 and EGFP. Representative micrographs of fluorescently labeled fibers from two replicates (n > 50 fibers). Scale bar 20 µm.

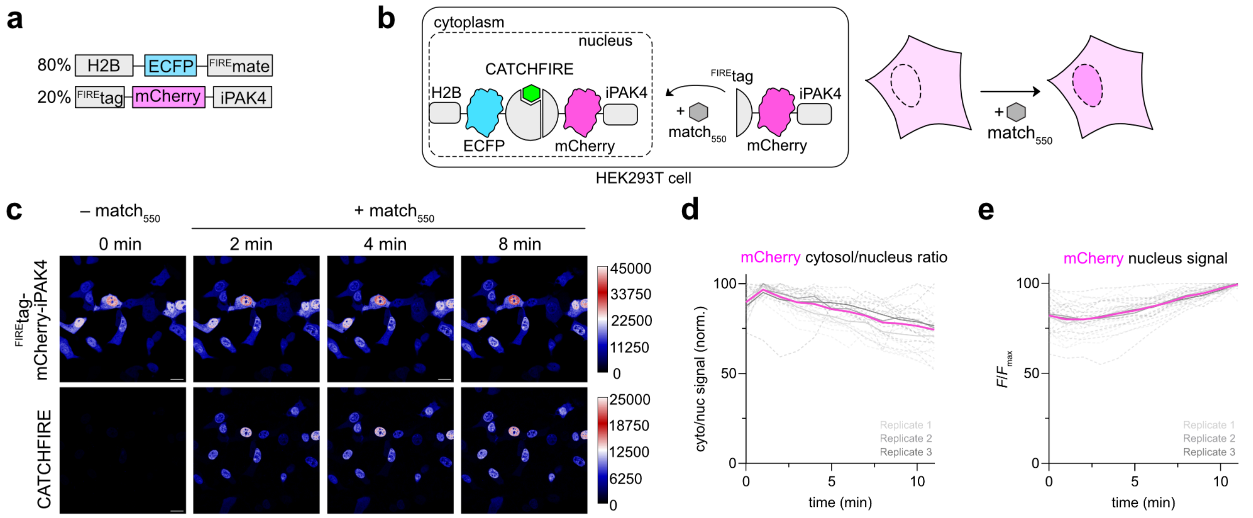


**Supplementary Figure 6. CATCHFIRE enables to control the localization of timestamp monomers.** **a** HEK293T were transfected with plasmids encoding H2B-ECFP-$^{FIRE}$mate (80%) and $^{FIRE}$tag-mCherry-iPAK4. **b** The day following transfection, cells were treated with 10 µM of $match_{550}$ and imaged by confocal time-lapse microscopy (frame rate = 1 frame / min). **c** Representative micrographs before and after $match_{550}$ addition in the red and green channel are shown. Scale bar 20 µm. **d,e** Temporal evolution of the ratio of mCherry fluorescence between the cytosol and nucleus (**d**) and the mCherry fluorescence intensity in the nucleus (**e**) of n = 28 cells from three replicates. The grey dashed lines correspond to the traces from individual cells, color-coded by their corresponding replicate. The grey full lines correspond to the mean of each replicate. The pink full line corresponds to the mean of the means of the three replicates.

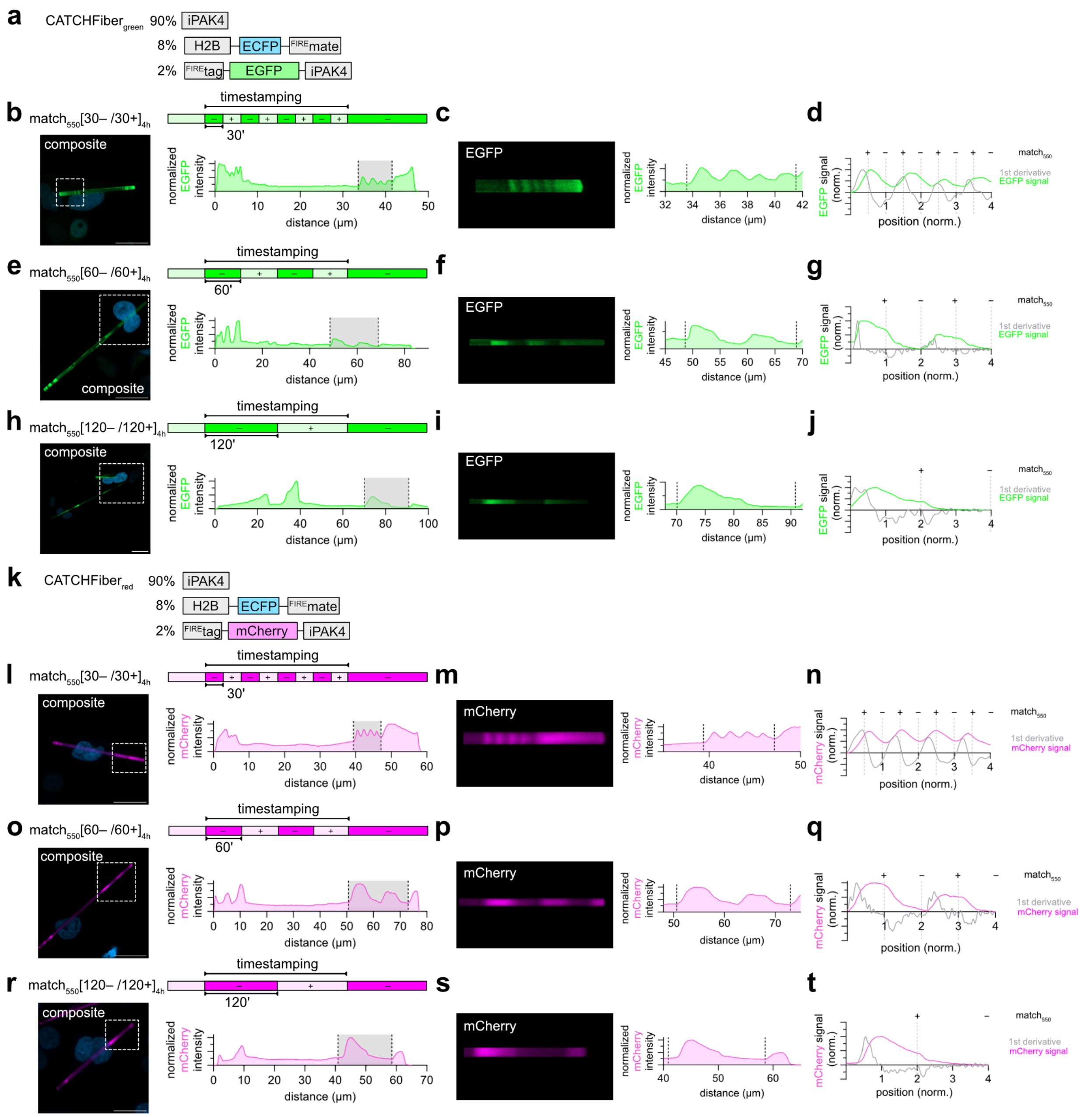


**Supplementary Figure 7. Evaluation of different pulse durations using CATCHFiber$_{green}$ and CATCHFiber$_{red}$ a-j** CATCHFiber$_{green}$ evaluation. **k-t** CATCHFiber$_{red}$ evaluation. **a,k** Plasmids used. **b-j,l-t** HEK293T cells expressing CATCHFiber$_{green}$ (**b-j**) or CATCHFiber$_{red}$ (**l-t**) were subjected to timestamping sequence match$_{550}$[30− / 30+]$_{4h}$ (**b-d, l-n**), match$_{550}$[60− / 60+]$_{4h}$ (**e-g, o-q**) and match$_{550}$ [120− / 120+]$_{4h}$ (**h-j, r-t**). For each pulse sequence, a representative micrograph as well as the fluorescence intensity plot (smoothed) along the displayed fiber is shown (**b,e,h,l,o,r**). The timestamping period is shown as a grey highlight in the full fluorescence intensity plot. A zoomed-in image of the timestamping patterns and the corresponding intensity plot (smoothed) is also shown (**c,f,i,m,p,s**). The dashed black lines positions were normalized to 0 and 4. The graphs in (**d,g,j,n,q,t**) show the fluorescence intensity (color) and the first derivative (gray) in function of the normalized position. The results are representative of n > 10 fibers (CATCHFiber$_{green}$) and n > 9 fibers (CATCHFiber$_{red}$) from two replicates. (**b,e,h,l,o,r**) Scale bar 20 µm.

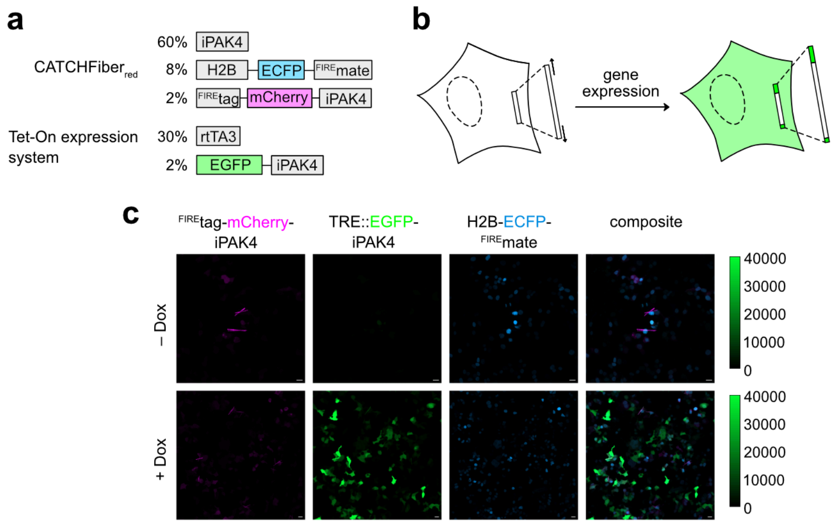


**Supplementary Figure 8. Doxycycline-induced expression of EGFP-iPAK4. a** Plasmids used for this study. **b** Schematic representation of HEK293T cell expressing CATCHFiber$_{red}$ and Tet-On expression system response to doxycycline addition, inducing EGFP expression. **c** Micrographs showing HEK293T cells expressing CATCHFiber$_{red}$ and Tet-On expression system, and subjected to the timestamping sequence match$_{500}$[30− / 60+]$_{4.5h}$. The cells were then imaged in the red, green and blue channels in the absence (above) or in the presence (below) of 2 µg/mL of doxycycline. The calibration bar in green shows the pixel intensity in the green channel. Representative micrographs from 6 fields of view for each condition from one replicate. Scale bar 20 µm

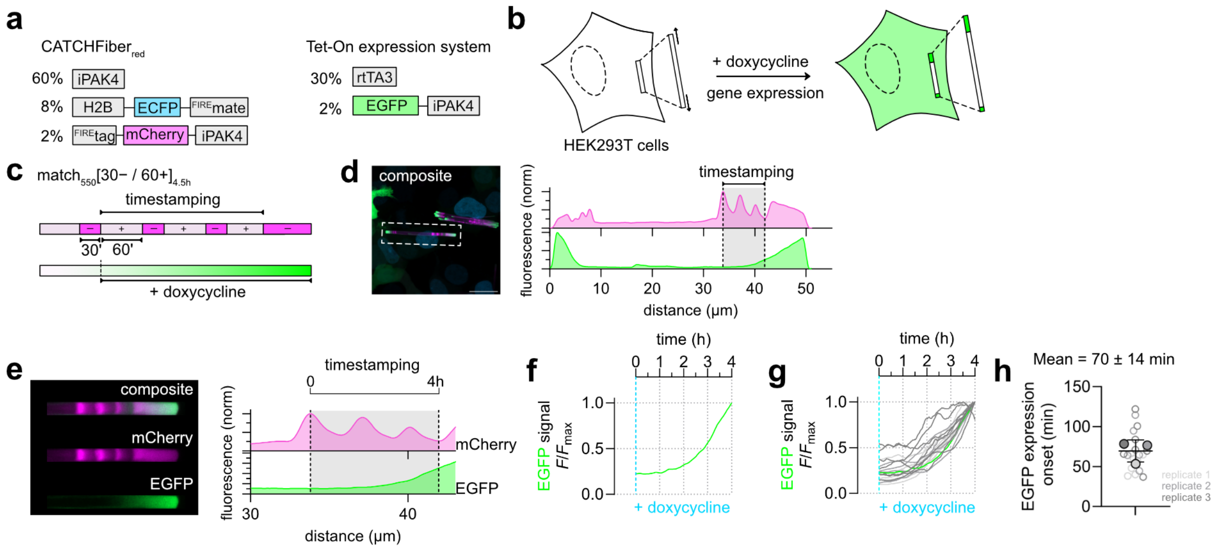


**Supplementary Figure 9. CATCHFiber enables the recording of gene expression under an inducible promoter. a** Plasmids used for this study. **b-h** HEK293T cells expressing CATCHFiber$_{red}$ and Tet-On expression system were subjected to the timestamping sequence match$_{550}$[30− / 60+]$_{4.5h}$ and doxycycline (2 µg/mL) was added along with the first match$_{550}$ addition (**c**). **d** Representative micrograph as well as the fluorescence intensity plot (smoothed) along the displayed fiber is shown. Scale bar 20 µm. The timestamping period is shown as a grey highlight in the full fluorescence intensity plot. **e** Image of the tip of the fast-growing edge of the fiber. The graph shows the intensity profile (smoothed) along the fiber. The dashed black lines positions were normalized to 0 and 4. The results are representative of n = 19 fibers from three replicates. **f** Temporal evolution of EGFP intensity (green, normalized) after temporal rescaling (see methods for detailed calculations). The blue dashed line indicates doxycycline addition time. **g** Temporal evolution of EGFP intensity from all 19 fibers imaged in this experiment (grey, green) after temporal rescaling. Each curve is color-coded according to the replicate the fiber comes from. In green is the temporal evolution of the fiber shown in (**d-f**). **h** EGFP expression onset times, identified from the curves in (**g**). On the graph, each fiber is color-coded according to the biological replicate it came from. The solid circles correspond to the mean of each biological replicate. The black line represents the mean ± SD of the three replicates (n = 19 fibers).

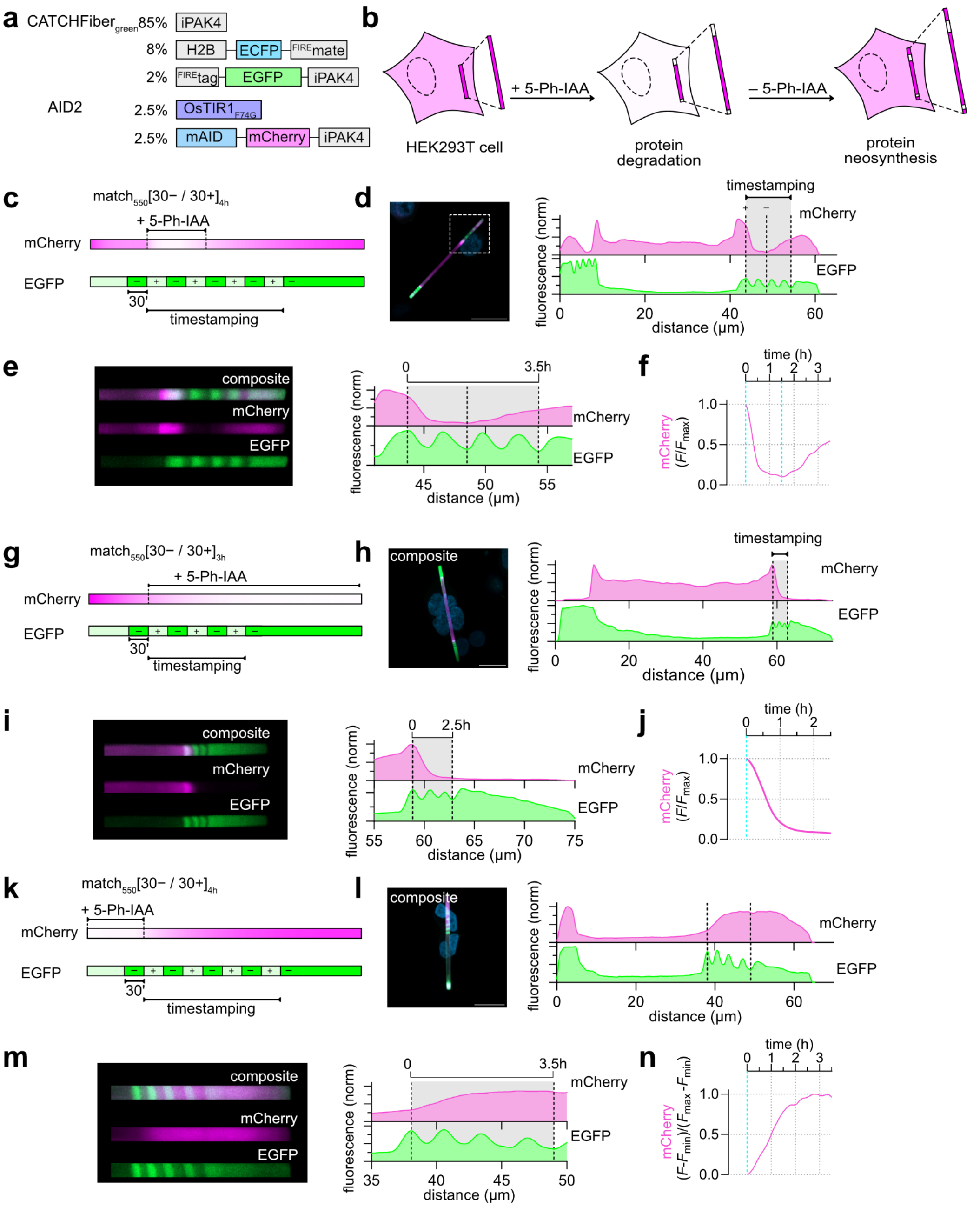

**Supplementary Figure 10. CATCHFiber enables the recording of proteasomal protein degradation and neosynthesis (full fiber information of Figure 3). a** Protein degradation and neosynthesis recording was evaluated through transfecting HEK293T cells with both CATCHFiber$_{green}$ and AID2 plasmids. **b** Proteasomal protein degradation was induced by addition of 5-Phenyl-indole-3-acetic acid (5-Ph-IAA, 1 µM), and neosynthesis was enabled through washing out 5-Ph-IAA. **c-f** Evaluation of the reversibility of the AID2 system using CATCHFiber. HEK293T cells expressing CATCHFiber$_{green}$ and AID2 were subjected to the timestamping sequence match$_{550}$ [30− / 30+]$_{4h}$. A composite image of a representative fiber as well as the corresponding intensity plot (smoothed) are shown in (**d**). A zoomed-in image of the end of the representative fiber and the corresponding intensity plot (smoothed) are shown in (**e**). The dashed black lines positions were normalized to 0 and 3.5 h. Degradation was induced through addition of 5-Ph-IAA along with the first match$_{550}$ addition, and neosynthesis was induced through washing out 5-Ph-IAA 1.5 h later. **f** Temporal evolution of mCherry intensity after temporal rescaling (see **Supplementary Text 1** for detailed calculations). The blue dashed lines indicate the duration of incubation in the presence of 5-Ph-IAA (1 µM). The results are representative of n = 33 fibers from two replicates. **g-j** Evaluation of the AID2-induced protein degradation dynamics. HEK293T cells expressing $CATCHFiber_{green}$ and AID2 were subjected to the timestamping sequence $match_{550}$ $[30- / 30+]_{3h}$. Degradation was induced through addition of 5-Ph-IAA along with the first $match_{550}$ addition. A composite image of a representative fiber as well as the corresponding intensity plot (smoothed) are shown in (**h**). A zoomed-in image of the end of the representative fiber and the corresponding intensity plot (smoothed) are shown in (**i**). The dashed black lines positions were normalized to 0 and 2.5 h. **j** Temporal evolution of mCherry intensity after temporal rescaling (see methods for detailed calculations). The blue dashed line indicates the addition of 5-Ph-IAA (1 µM). The results are representative of n = 31 fibers from four replicates **k-n** Evaluation of the neosynthesis dynamics following 5-Ph-IAA washout. HEK293T cells expressing $CATCHFiber_{green}$ and AID2 pretreated with 5-Ph-IAA were subjected to the timestamping sequence $match_{550}$ $[30- / 30+]_{4h}$. Neosynthesis was induced through washing out of 5-Ph-IAA along with the first $match_{550}$ addition. A composite image of a representative fiber as well as the corresponding intensity plot (smoothed) are shown in (**l**). A zoomed-in image of the end of the representative fiber and the corresponding intensity plot (smoothed) are shown in (**m**). The dashed black lines positions were normalized to 0 and 3.5 h. **n** Temporal evolution of mCherry intensity after temporal rescaling (see **Supplementary Text 1** for detailed calculations). The blue dashed line indicates the washing out of 5-Ph-IAA (1 µM). The results are representative of n = 29 fibers from four replicates.

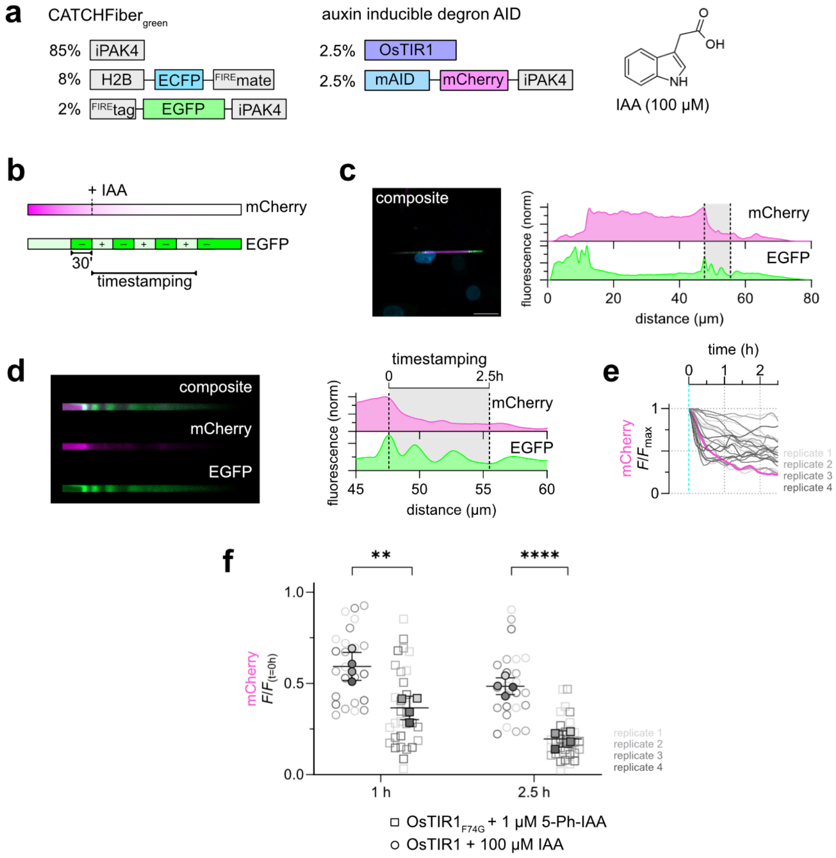


**Supplementary Figure 11. CATCHFiber enables the kinetics comparison between AID and AID2 for proteasomal protein degradation.** Protein degradation using AID induced in HEK293T cells through expression of $\text{CATCHFiber}_{\text{green}}$ and auxin-inducible degron AID plasmids and addition of indole-3-acetic acid (IAA) (**a**). HEK293T cells expressing $\text{CATCHFiber}_{\text{green}}$ and AID2 were subjected to the timestamping sequence $\text{match}_{550}[30- / 30+]_{3h}$. Degradation was induced through addition of 5-Ph-IAA along with the first $\text{match}_{550}$ addition (**b**). A composite image of a representative fiber as well as the corresponding full intensity plot (smoothed) are shown in (**c**). Scale bar 20 µm. A zoomed-in image of the end of the representative fiber and the corresponding intensity plot (smoothed) are shown in (**d**). The dashed black lines positions were normalized to 0 and 2.5 h. The results are representative of n = 24 fibers from four replicates **e** Temporal evolution of normalized mCherry intensity from all 24 fibers imaged in this experiment after temporal rescaling. Each curve is color-coded according to the replicate the fiber comes from. In pink is the temporal evolution of the fiber shown in **d** **f** Graph showing the evaluation of the degradation extent of AID2 (squares, see **Figure 3** for full analysis) vs AID (circles) at t = 1 h and t = 2.5 h of timestamping measured as the normalized intensity of mCherry signal. On the graph each fiber is color-coded according to the biological replicate it came from. The solid squares and circles correspond to the mean of each biological replicate. The black line represents the mean ± SD of the four replicates (n = 31 fibers for AID2 and n = 24 fibers for AID from four biological replicates). An unpaired two-tailed t-test assuming equal variance was used to compare the distributions for AID2 and AID at t = 1 h (**$P$ = 0.0039) and t = 2.5 h (****$P$ < 0.0001).

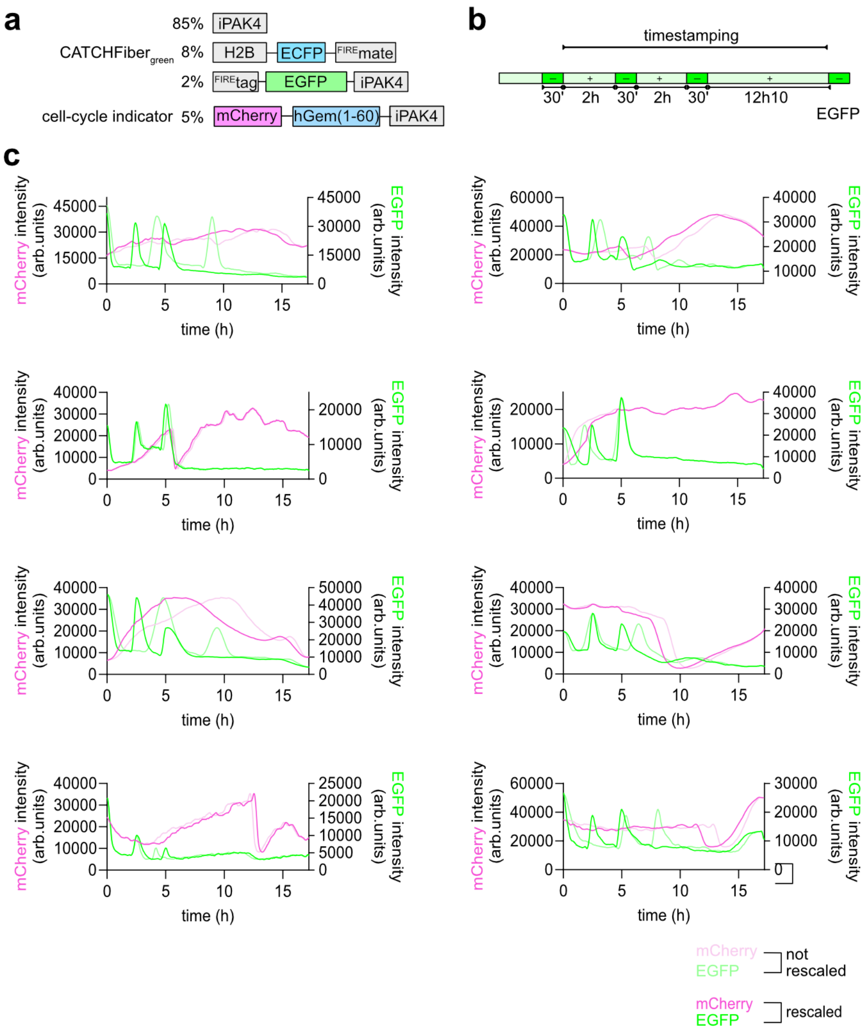


**Supplementary Figure 12. CATCHFiber-based timestamps helps the identification of fibers which show stalled growth during long-term recording of cell-cycle dynamics.** HEK293T cells expressing CATCHFiber$_{green}$ and the cell cycle indicator (**a**) were submitted to the timestamping sequence indicated in (**b**). **c** Temporal evolution of mCherry and EGFP signals during the timestamping period, using all timestamps for temporal rescaling (dark colors) and using only the first and last timestamp information (light colors).

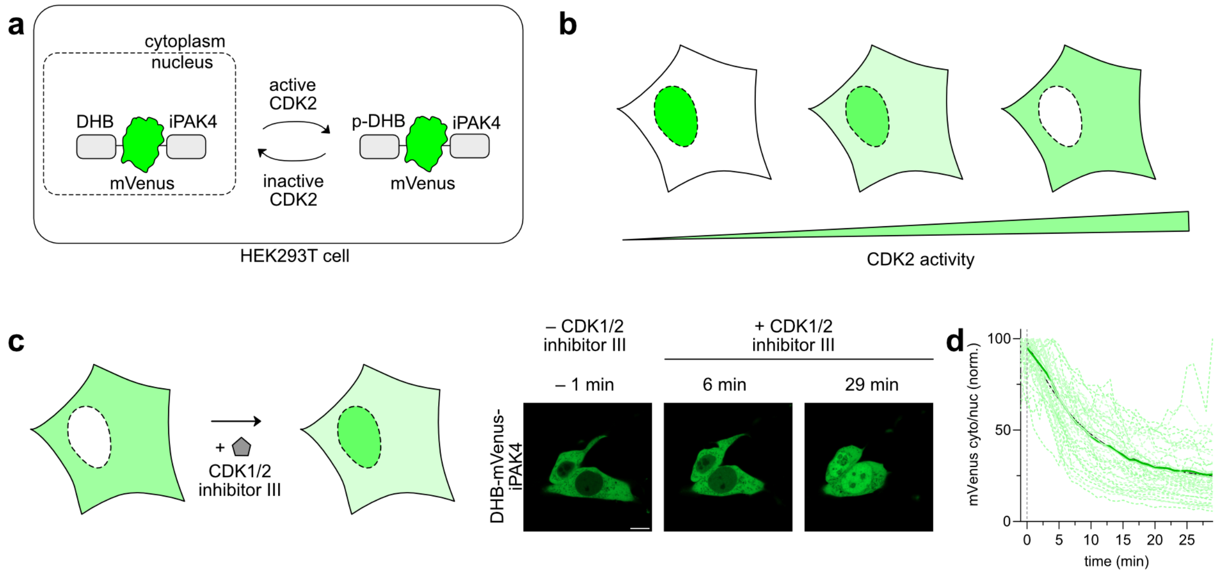


**Supplementary Figure 13. DHB-mVenus-iPAK4 reports on CDK2 activity. a,b** DHB changes its localization in function of CDK2 activity. Upon phosphorylation, it translocates from the nucleus to the cytosol. Evaluation of its localization gives access to CDK2 activity. **c** HEK293T were transfected with plasmids encoding DHB-mVenus-iPAK4. The day following transfection, cells were treated with 10 µM of CDK1/2 inhibitor **III** and imaged by confocal time-lapse microscopy (frame rate = 1 frame / min). Representative micrographs before and after CDK1/2 inhibitor **III** addition in the green channel are shown. Scale bar 20 µm. **d** Temporal evolution of the ratio of mVenus fluorescence between the cytosol and nucleus of n = 43 cells from two replicates. The green dashed lines correspond to the traces from individual cells, color coded by their corresponding replicate. The green full lines correspond to the mean of each replicate. The green bold line corresponds to the mean of the means of the two replicates, and the black line to the corresponding fit with a one-phase decay model.

**Annex Figures 1 and 2– Data extraction and analysis workflow**

**Data extraction workflow**

1. Each acquisition is a multichannel z-stack of n slices required to capture the whole tilted fiber. In case of a fiber fully captured within a single plane, the starting point is a multichannel snap.

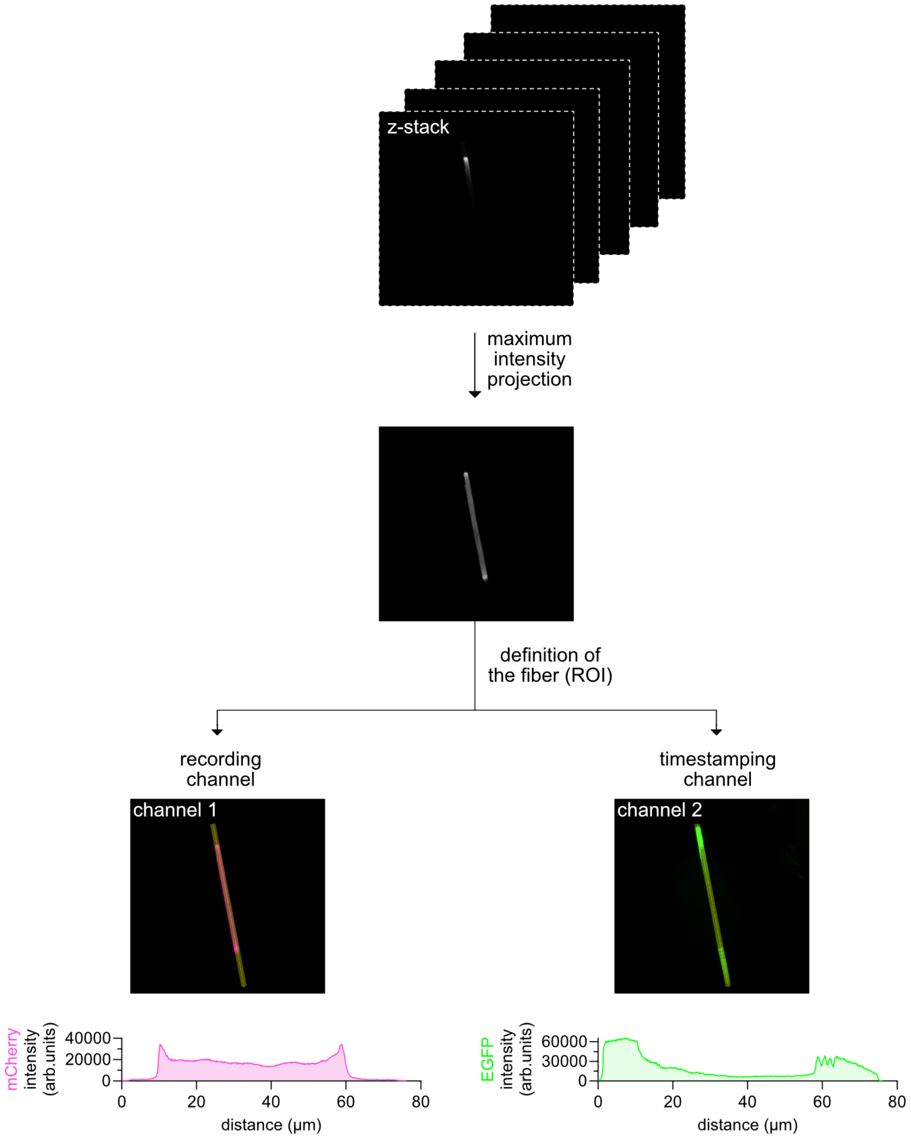


2. On the maximum intensity projection, for each channel, a line ROI around the fiber is drawn, the fluorescence intensity profile extracted and saved.

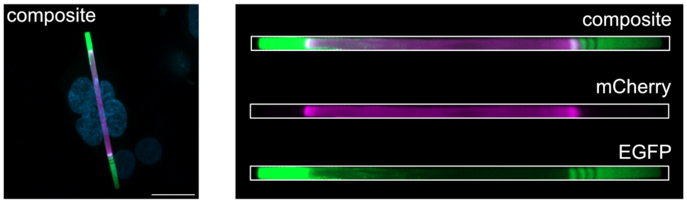


3. For each fiber, the maximum intensity projection as well as the straightened line ROI is saved. Composites of the projections and straightened fibers are generated for the displayed fibers

**Data analysis and formatting workflow**

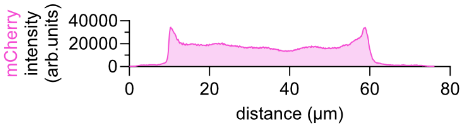


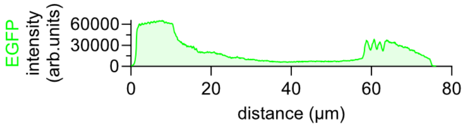


smoothing and normalization

1. The fluorescent intensity profiles are plotted for each channel, normalized to $F_{max}$ and smoothed.

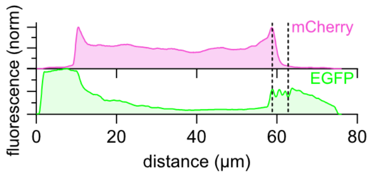


calculation of first derivative and identification of cancellation points

2. The first derivative of the intensity profile of the fiber in the timestamping channel is plotted. The additions and washouts of $match_{550}$ are identified as cancellation points of the first derivative. To each cancellation point, the fiducial timestamp (**ground truth time**) is attributed.

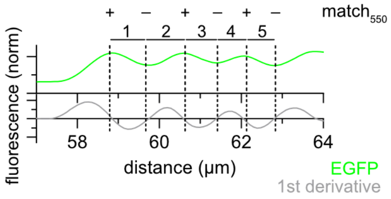


| distance (µm) | 58.8 | 59.7 | 60.6 | 61.4 | 62.1 | 62.8 |
|---|---|---|---|---|---|---|
| **ground truth time (h)** | 0 | 0.5 | 1 | 1.5 | 2 | 2.5 |

| segment | 1 | 2 | 3 | 4 | 5 |
|---|---|---|---|---|---|
| **length (µm)** | 0.9 | 0.9 | 0.8 | 0.7 | 0.7 |

identification of timestamping borders

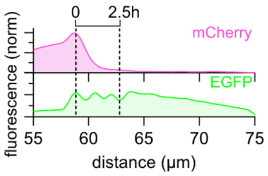


re-normalization between timestamps

3. The distance axis is normalized for each segment bounded by timestamps (**rescaled curves**). In the example shown here, the fiber grew 4 µm in 2.5 h. In the hypothesis of linear growth, the distance between two fiducial timestamps should be of 0.8 µm. In this particular example, the linear growth assumption is met as the length of the segments is very close to 0.8 µm, and the difference between the rescaled and the not rescaled curves is small.

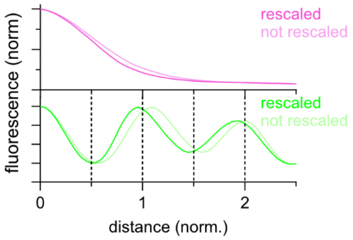

**Supplementary Table 1 – Plasmids ratios**

| Figure | Plasmid ratio |
|---|---|
| **Fig 2.** | FLAG-iPAK4 (pAG1903, 85%), H2B-ECFP-$^{\text{FIRE}}$mate (pAG1247, 8%), $^{\text{FIRE}}$tag-mCherry-iPAK4 (pAG1910, 2%), EGFP-iPAK4 (pAG1904, 5%) |
| **Fig 3.** | FLAG-iPAK4 (pAG1903, 85%), H2B-ECFP-$^{\text{FIRE}}$mate (pAG1247, 8%), $^{\text{FIRE}}$tag-EGFP-iPAK4 (pAG1909, 2%), OsTIR1$_{\text{F74G}}$ (pAG2063, 2.5%), mAID-mCherry-iPAK4 (pAG2055, 2.5%) |
| **Fig 4.** | FLAG-iPAK4 (pAG1903, 85%), H2B-ECFP-$^{\text{FIRE}}$mate (pAG1247, 8%), $^{\text{FIRE}}$tag-EGFP-iPAK4 (pAG1909, 2%), OsTIR1$_{\text{F74G}}$ (pAG2063, 2.5%), mAID-mCherry-iPAK4 (pAG2055, 2.5%) |
| **Fig 5.** | FLAG-iPAK4 (pAG1903, 85%), H2B-ECFP-$^{\text{FIRE}}$mate (pAG1247, 8%), $^{\text{FIRE}}$tag-EGFP-iPAK4 (pAG1909, 2%), mCherry-hGem$_{1\text{-}60}$-iPAK4 (pAG2101, 5%) |
| **Fig 6.** | FLAG-iPAK4 (pAG1903, 85%), H2B-ECFP-$^{\text{FIRE}}$mate (pAG1247, 8%), $^{\text{FIRE}}$tag-mCherry-iPAK4 (pAG1910, 2%), DHB-mVenus-iPAK4 (pAG2099, 5%) |
| **SI Fig 1.** | (1) FLAG-iPAK4 (pAG1903, 90%), $^{\text{FIRE}}$tag-EGFP-iPAK4 (pAG1909, 10%)<br>(2) FLAG-iPAK4 (pAG1903, 90%), $^{\text{FIRE}}$tag-mCherry-iPAK4 (pAG1910, 10%)<br>(3) FLAG-iPAK4 (pAG1903, 90%), $^{\text{FIRE}}$tag-emiRFP670-iPAK4 (pAG1911, 10%)<br>(4) FLAG-iPAK4 (pAG1903, 90%), EGFP-iPAK4 (pAG1904, 10%)<br>(5) FLAG-iPAK4 (pAG1903, 90%), mCherry-iPAK4 (pAG1907, 10%)<br>(6) FLAG-iPAK4 (pAG1903, 90%), emiRFP670-iPAK4 (pAG1908, 10%) |
| **SI Fig 2.** | (1) FLAG-iPAK4 (pAG1903, 90%), $^{\text{FIRE}}$tag-EGFP-iPAK4 (pAG1909, 10%)<br>(2) FLAG-iPAK4 (pAG1903, 90%), $^{\text{FIRE}}$tag-mCherry-iPAK4 (pAG1910, 10%)<br>(3) FLAG-iPAK4 (pAG1903, 90%), $^{\text{FIRE}}$tag-emiRFP670-iPAK4 (pAG1911, 10%) |
| **SI Fig 3.** | (1) FLAG-iPAK4 (pAG1903, 90%), EGFP-iPAK4 (pAG1904, 10%)<br>(2) FLAG-iPAK4 (pAG1903, 90%), mCherry-iPAK4 (pAG1907, 10%)<br>(3) FLAG-iPAK4 (pAG1903, 90%), emiRFP670-iPAK4 (pAG1908, 10%) |
| **SI Fig 4.** | (1) FLAG-iPAK4 (pAG1903, 90%), EGFP-iPAK4 (pAG1904, 5%), mCherry-iPAK4 (pAG1907, 5%)<br>(2) FLAG-iPAK4 (pAG1903, 90%), emiRFP670-iPAK4 (pAG1908, 5%), mCherry-iPAK4 (pAG1907, 5%)<br>(3) FLAG-iPAK4 (pAG1903, 90%), emiRFP670-iPAK4 (pAG1908, 5%), EGFP-iPAK4 (pAG1904, 5%) |
| **SI Fig 5.** | (1) FLAG-iPAK4 (pAG1903, 90%), EGFP-iPAK4 (pAG1904, 5%), mCherry-iPAK4 (pAG1907, 5%)<br>(2) FLAG-iPAK4 (pAG1903, 90%), emiRFP670-iPAK4 (pAG1908, 5%), EGFP-iPAK4 (pAG1904, 5%)<br>(3) FLAG-iPAK4 (pAG1903, 90%), emiRFP670-iPAK4 (pAG1908, 5%), mCherry-iPAK4 (pAG1907, 5%) |
| **SI Fig 6.** | H2B-ECFP-$^{\text{FIRE}}$mate (pAG1247, 80%), $^{\text{FIRE}}$tag-mCherry-iPAK4 (pAG1910, 20%) |
| **SI Fig 7.** | (1) FLAG-iPAK4 (pAG1903, 90%), H2B-ECFP-$^{\text{FIRE}}$mate (pAG1247, 8%), $^{\text{FIRE}}$tag-EGFP-iPAK4 (pAG1909, 2%)<br>(2) FLAG-iPAK4 (pAG1903, 90%), H2B-ECFP-$^{\text{FIRE}}$mate (pAG1247, 8%), $^{\text{FIRE}}$tag-mCherry-iPAK4 (pAG1910, 2%) |
| **SI Fig 8.** | FLAG-iPAK4 (pAG1903, 60%), H2B-ECFP-$^{\text{FIRE}}$mate (pAG1247, 8%), $^{\text{FIRE}}$tag-mCherry-iPAK4 (pAG1910, 2%), rtTA3 (pAG2064, 30%), TRE::EGFP-iPAK4 (Addgene plasmid # 187446, 2%) |

| | |
|---|---|
| **SI Fig 9.** | FLAG-iPAK4 (pAG1903, 60%), H2B-ECFP-$^{FIRE}$mate (pAG1247, 8%), $^{FIRE}$tag-mCherry-iPAK4 (pAG1910, 2%), rtTA3 (pAG2064, 30%), TRE::EGFP-iPAK4 (Addgene plasmid # 187446, 2%) |
| **SI Fig 11.** | FLAG-iPAK4 (pAG1903, 85%), H2B-ECFP-$^{FIRE}$mate (pAG1247, 8%), $^{FIRE}$tag-EGFP-iPAK4 (pAG1909, 2%), OsTIR1$_{F74G}$ (pAG2063, 2.5%), mAID-mCherry-iPAK4 (pAG2055, 2.5%) |
| **SI Fig 12.** | AID2: FLAG-iPAK4 (pAG1903, 85%), H2B-ECFP-$^{FIRE}$mate (pAG1247, 8%), $^{FIRE}$tag-EGFP-iPAK4 (pAG1909, 2%), OsTIR1$_{F74G}$ (pAG2063, 2.5%), mAID-mCherry-iPAK4 (pAG2055, 2.5%)<br>AID: FLAG-iPAK4 (pAG1903, 85%), H2B-ECFP-$^{FIRE}$mate (pAG1247, 8%), $^{FIRE}$tag-EGFP-iPAK4 (pAG1909, 2%), OsTIR1 (pAG2062, 2.5%), mAID-mCherry-iPAK4 (pAG2055, 2.5%) |
| **SI Fig 13.** | FLAG-iPAK4 (pAG1903, 85%), H2B-ECFP-$^{FIRE}$mate (pAG1247, 8%), $^{FIRE}$tag-EGFP-iPAK4 (pAG1909, 2%), mCherry-hGem$_{1-60}$-iPAK4 (pAG2101, 5%) |
| **SI Fig 14.** | DHB-mVenus-iPAK4, pAG2099 (100%) |

### Supplementary Table 2 – Plasmids used in this study

| Plasmid number | ORF | Sequence |
|---|---|---|
| pAG1247 (already published[2]) | H2B-ECFP-FIREmate | atgcccgaacctgcgaagtcagcgcccgctcccaaaaaggctctaaaaagctgtcgccaagaccca<br>gaagaaggggataagaaaaggcgtaagaccaggaaagagagttacgccatttacgtgtacaaagta<br>ctaaaacaagtccacccggacactggcatctcctcaaaggcgatgggcattatgaactcatttgtaaacga<br>catcttcgagcgcatcgccggagaagcgtcgcgcctggcgcattacaacaagcgctccactatcacatcc<br>cgggagatccagacggccgtgcgcctgctcctgcccggagaactggccaaacacgctgtgtctgagggc<br>acaaaggccgtgaccaagtacaccagctccaagggtagtgctggtggtatggtgagcaagggcgagga<br>gctgttcaccggggtggtgcccatcctggtcgagctggacggcgacgtaaacggccacaagttcagcgtg<br>tccggcgagggcgagggcgatgccacctacggcaagctgaccctgaagttcatctgcaccaccggcaa<br>gctgcccgtgccctggcccaccctcgtgaccaccctgacctggggcgtgcagtgcttcagccgctaccccg<br>accacatgaagcagcacgacttcttcaagtccgccatgcccgaaggctacgtccaggagcgcaccatctt<br>cttcaaggacgacggcaactacaagacccgcgccgaggtgaagttcgagggcgacaccctggtgaac<br>cgcatcgagctgaagggcatcgacttcaaggaggacggcaacatcctggggcacaagctggagtacaa<br>ctacatcagccacaacgtctatatcaccgccgacaagcagaagaacggcatcaaggccaacttcaagat<br>ccgccacaacatcgaggacggcagcgtgcagctcgccgaccactaccagcagaacacccccatcggc<br>gacggccccgtgctgctgcccgacaaccactacctgagcacccagtccgccctgagcaaagaccccaa<br>cgagaagcgcgatcacatggtcctgctggagttcgtgaccgccgccgggatcactctcggcatggacga<br>gctgtacaagggagcaagtggaatggagcatgttgcctttggcagtgaggacatcgagaacactctggcc<br>aatatggacgacgaacaactggataggttggcctttggcgtaattcagctcgatggtgacgggaatatcctg<br>ctgtacaatgctgctgaaggggacatcactggcagagatcccaaacaggtgattgggaagaacttcttca<br>aggatgttgcacctggaacggatactcccgagttttacggcaaattcaaggaaggcgcagcgtcaggga<br>atctgaacaccatgttcgaatggacgataccgacaagcaggggaccaaccaaggtcaaggtgcacttga<br>agaaagccctttcc |
| pAG1903 | FLAG-SGS-INKAbox-Pak4cat | atggactacaaggacgacgatgataagtccggatccgaagcagaggactggacagcagccctactga<br>acaggggtcgcagtcgccagcccctggtactaggggacaattgctttgctgacttggtgcacaactggatg<br>gagctgcctgaggaattccccgccgcccctgctgttcctgggccccctggcccccgctcaccacagcggg<br>agccacagcgagtatcccatgagcagttccgggctgccctgcagctggtggtggacccaggcgaccccc<br>gctcctacctggacaacttcatcaagattggcgagggctccacgggcatcgtgtgcatcgccaccgtgcgc<br>agctcgggcaagctggtggccgtcaagaagatggacctgcgcaagcagcagaggcgcgagctgctctt<br>caacgaggtggtaatcatgagggactaccagcacgagaatgtggtggagatgtacaacagctacctggt<br>gggggacgagctctgggtggtcatggagttcctggaaggaggcgccctcaccgacatcgtcacccacac<br>caggatgaacgaggagcagatcgcagccgtgtgccttgcagtgctgcaggccctgtcggtgctccacgcc<br>cagggcgtcatccaccgggacatcaagagcgactcgatcctgctgacccatgatggcagggtgaagctg<br>tcagactttgggttctgcgcccaggtgagcaaggaagtgccccgaaggaagtcgctggtcggcacgccct<br>actggatggccccagagctcatctcccgccttccctacgggccagaggtagacatctggtcgctggggata<br>atggtgattgagatggtggacggagagccccctacttcaacgagccacccctcaaagccatgaagatg<br>attcgggacaacctgccaccccgactgaagaacctgcacaaggtgtcgccatccctgaagggcttcctgg<br>accgcctgctggtgcgagaccctgcccagcgggccacggcagccgagctgctgaagcacccattcctgg<br>ccaaggcagggccgcctgccagcatcgtgcccctcatgcgccagaaccgcaccaga |
| pAG1904 | EGFP-SGGS-INKAbox-Pak4cat | atgagtaaaggagaagaacttttcactggagttgtcccaattcttgttgaattagatggtgatgttaatgggtac<br>aaattttctgtcagtggagagggtgaaggtgatgcaacatacggaaaacttacccttaaatttatttgcactac<br>tggaaaactacctgttccatggccaacacttgtcactactctcacttatggtgttcaatgcttttcaagatatcca<br>gatcatatgaagcggcacgacttcttcaagagcgccatgcctgagggatacgtgcaggagaggaccatct<br>tcttcaaggacgacgggaactacaagacacgtgctgaagtcaagtttgagggagacaccctcgtcaaca<br>ggatcgagcttaagggaatcgatttcaaggaggacggaaacatcctcggccacaagttggaatacaact<br>acaactcccacaacgtatacatcatggccgacaagcaaaagaacggcatcaaagccaacttcaagacc<br>cgccacaacatcgaagacggcggcgtgcaactcgctgatcattatcaacaaaatactccaattggcgatg<br>accctgtccttttaccagacaaccattacctgtccacacaatctgccctttcgaaagatcccaacgaaaaga<br>gagaccacatggtcattcttgagtttgtaacggctgctgggattacacatggcatggatgaactatacaaatc<br>aggtggatccgaagcagaggactggacagcagccctactgaacaggggtcgcagtcgccagcccctg<br>gtactaggggacaattgctttgctgacttggtgcacaactggatggagctgcctgaggaattccccgccgcc |

| | | |
|---|---|---|
| | | cctgctgttcctgggcccccctggcccccgctcaccacagcgggagccacagcgagtatcccatgagcagt<br>tccgggctgccctgcagctggtggtggacccaggcgacccccgctcctacctggacaacttcatcaagatt<br>ggcgagggctccacgggcatcgtgtgcatcgccaccgtgcgcagctcgggcaagctggtggccgtcaag<br>aagatggacctgcgcaagcagcagaggcgcgagctgctcttcaacgaggtggtaatcatgagggacta<br>ccagcacgagaatgtggtggagatgtacaacagctacctggtgggggacgagctctgggtggtcatgga<br>gttcctggaaggaggcgccctcaccgacatcgtcacccacaccaggatgaacgaggagcagatcgcag<br>ccgtgtgccttgcagtgctgcaggccctgtcggtgctccacgcccagggcgtcatccaccgggacatcaa<br>gagcgactcgatcctgctgacccatgatggcagggtgaagctgtcagactttgggttctgcgcccaggtga<br>gcaaggaagtgccccgaaggaagtcgctggtcggcacgccctactggatggccccagagctcatctccc<br>gccttccctacgggccagaggtagacatctggtcgctggggataatggtgattgagatggtggacggaga<br>gccccccctacttcaacgagccacccctcaaagccatgaagatgattcgggacaacctgccaccccgact<br>gaagaacctgcacaaggtgtcgccatccctgaagggcttcctggaccgcctgctggtgcgagaccctgcc<br>cagcgggccacggcagccgagctgctgaagcacccattcctggccaaggcagggccgcctgccagca<br>tcgtgcccctcatgcgccagaaccgcaccaga |
| pAG1907 | mCherry-SGGS-INKAbox-Pak4cat | Atggtgagcaagggcgaggaggataacatggccatcatcaaggagttcatgcgcttcaaggtgcacatg<br>gagggctccgtgaacggccacgagttcgagatcgagggcgagggcgagggccgcccctacgagggc<br>acccagaccgccaagctgaaggtgaccaagggtggccccctgcccttcgcctgggacatcctgtcccctc<br>agttcatgtacggctccaaggcctacgtgaagcaccccgccgacatccccgactacttgaagctgtccttcc<br>ccgagggcttcaagtgggagcgcgtgatgaacttcgaggacggcggcgtggtgaccgtgacccaggact<br>cctccctgcaggacggcgagttcatctacaaggtgaagctgcgcggcaccaacttcccctccgacggccc<br>cgtaatgcagaagaagaccatgggctgggaggcctcctccgagcggatgtaccccgaggacggcgcc<br>ctgaagggcgagatcaagcagaggctgaagctgaaggacggcggccactacgacgctgaggtcaag<br>accacctacaaggccaagaagcccgtgcagctgcccggcgcctacaacgtcaacatcaagttggacat<br>cacctcccacaacgaggactacaccatcgtggaacagtacgaacgcgccgagggccgccactccacc<br>ggcggcatggacgagctgtacaagtcaggtggatccgaagcagaggactggacagcagccctactga<br>acaggggtcgcagtcgccagcccctggtactaggggacaattgctttgctgacttggtgcacaactggatg<br>gagctgcctgaggaattccccgccgcccctgctgttcctgggcccccctggcccccgctcaccacagcggg<br>agccacagcgagtatcccatgagcagttccgggctgccctgcagctggtggtggacccaggcgaccccc<br>gctcctacctggacaacttcatcaagattggcgagggctccacgggcatcgtgtgcatcgccaccgtgcgc<br>agctcgggcaagctggtggccgtcaagaagatggacctgcgcaagcagcagaggcgcgagctgctctt<br>caacgaggtggtaatcatgagggactaccagcacgagaatgtggtggagatgtacaacagctacctggt<br>gggggacgagctctgggtggtcatggagttcctggaaggaggcgccctcaccgacatcgtcacccacac<br>caggatgaacgaggagcagatcgcagccgtgtgccttgcagtgctgcaggccctgtcggtgctccacgcc<br>cagggcgtcatccaccgggacatcaagagcgactcgatcctgctgacccatgatggcagggtgaagctg<br>tcagactttgggttctgcgcccaggtgagcaaggaagtgccccgaaggaagtcgctggtcggcacgccct<br>actggatggccccagagctcatctcccgccttccctacgggccagaggtagacatctggtcgctggggata<br>atggtgattgagatggtggacggagagccccccctacttcaacgagccacccctcaaagccatgaagatg<br>attcgggacaacctgccaccccgactgaagaacctgcacaaggtgtcgccatccctgaagggcttcctgg<br>accgcctgctggtgcgagaccctgcccagcgggccacggcagccgagctgctgaagcacccattcctgg<br>ccaaggcagggccgcctgccagcatcgtgcccctcatgcgccagaaccgcaccaga |
| pAG1908 | emiRFP670-SGGS-INKAbox-Pak4cat | Atggcggaaggatccgtcgccaggcagcctgacctcttgacctgcgaacatgaagagatccacctcgcc<br>ggctcgatccagccgcatggcgcgcttctggtcgtcagcgaacatgatcatcgcgtcatccaggccagcg<br>ccaacgccgcggaatttctgaatctcggaagcgtactcggcgttccgctcgccgagatcgacggcgatctg<br>ttgatcaagatcctgccgcatctcgatcccaccgccgaaggcatgccggtcgcggtgcgctgccggatcg<br>gcaatccctctacggagtactgcggtctgatgcatcggcctccggaaggcgggctgatcatcgaactcga<br>acgtgccggcccgtcgatcgatctgtcaggcacgctggcgccggcgctggagcggatccgcacggcgg<br>gttcactgcgcgcgctgtgcgatgacaccgtgctgctgtttcagcagtgcaccggctacgaccgggtgatg<br>gtgtatcgtttcgatgagcaaggccacggcctggtattctccgagtgccatgtgcctgggctcgaatcctattt<br>cggcaaccgctatccgtcgtcgactgtcccgcagatggcgcggcagctgtacgtgcggcagcgcgtccg<br>cgtgctggtcgacgtcacctatcagccggtgccgctggagccgcggctgtcgccgctgaccgggcgcgat<br>ctcgacatgtcgggctgcttcctgcgctcgatgtcgccgtgccatctgcagttcctgaaggacatgggcgtgc<br>gcgccaccctggcggtgtcgctggtggtcggcggcaagctgtggggcctggttgtctgtcaccattatctgcc<br>gcgcttcatccgtttcgagctgcgggcgatctgcaaacggctcgccgaaaggatcgcgacgcggatcacc<br>gcgcttgagagctcaggtggatccgaagcagaggactggacagcagccctactgaacaggggtcgcag<br>tcgccagcccctggtactaggggacaattgctttgctgacttggtgcacaactggatggagctgcctgagga<br>attccccgccgcccctgctgttcctgggcccccctggcccccgctcaccacagcgggagccacagcgagta |

| | | |
|---|---|---|
| | | tcccatgagcagttccgggctgccctgcagctggtggtggacccaggcgacccccgctcctacctggaca<br>acttcatcaagattggcgagggctccacgggcatcgtgtgcatcgccaccgtgcgcagctcgggcaagct<br>ggtggccgtcaagaagatggacctgcgcaagcagcagaggcgcgagctgctcttcaacgaggtggtaa<br>tcatgagggactaccagcacgagaatgtggtggagatgtacaacagctacctggtgggggacgagctct<br>gggtggtcatggagttcctggaaggaggcgccctcaccgacatcgtcacccacaccaggatgaacgag<br>gagcagatcgcagccgtgtgccttgcagtgctgcaggccctgtcggtgctccacgcccagggcgtcatcc<br>accgggacatcaagagcgactcgatcctgctgacccatgatggcagggtgaagctgtcagactttgggttc<br>tgcgcccaggtgagcaaggaagtgccccgaaggaagtcgctggtcggcacgccctactggatggcccc<br>agagctcatctcccgccttccctacgggccagaggtagacatctggtcgctggggataatggtgattgagat<br>ggtggacggagagcccccctacttcaacgagccacccctcaaagccatgaagatgattcgggacaacct<br>gccaccccgactgaagaacctgcacaaggtgtcgccatccctgaagggcttcctggaccgcctgctggtg<br>cgagaccctgcccagcgggccacggcagccgagctgctgaagcacccattcctggccaaggcagggc<br>cgcctgccagcatcgtgcccctcatgcgccagaaccgcaccaga |
| pAG1909 | FIREtag-SGGGGSGG-EGFP-SGGS-INKAbox-PAK4cat | atgggtgacagatattgggtctttgtgaaacgggtgagcggcgggggaggctccggagggagtaaagga<br>gaagaacttttcactggagttgtcccaattcttgttgaattagatggtgatgttaatgggtacaaattttctgtcag<br>tggagagggtgaaggtgatgcaacatacggaaaacttacccttaaatttatttgcactactggaaaactacc<br>tgttccatggccaacacttgtcactactctcacttatggtgttcaatgcttttcaagatatccagatcatatgaag<br>cggcacgacttcttcaagagcgccatgcctgagggatacgtgcaggagaggaccatcttcttcaaggacg<br>acgggaactacaagacacgtgctgaagtcaagtttgagggagacaccctcgtcaacaggatcgagctta<br>agggaatcgatttcaaggaggacggaaacatcctcggccacaagttggaatacaactacaactcccaca<br>acgtatacatcatggccgacaagcaaaagaacggcatcaaagccaacttcaagacccgccacaacatc<br>gaagacggcggcgtgcaactcgctgatcattatcaacaaaatactccaattggcgatgaccctgtcctttta<br>ccagacaaccattacctgtccacacaatctgccctttcgaaagatcccaacgaaaagagagaccacatg<br>gtcattcttgagtttgtaacggctgctgggattacacatggcatggatgaactatacaaatcaggtggatccg<br>aagcagaggactggacagcagccctactgaacaggggtcgcagtcgccagcccctggtactagggga<br>caattgctttgctgacttggtgcacaactggatggagctgcctgaggaattccccgccgcccctgctgttcctg<br>ggcccctggcccccgctcaccacagcgggagccacagcgagtatcccatgagcagttccgggctgcc<br>ctgcagctggtggtggacccaggcgacccccgctcctacctggacaacttcatcaagattggcgagggct<br>ccacgggcatcgtgtgcatcgccaccgtgcgcagctcgggcaagctggtggccgtcaagaagatggacc<br>tgcgcaagcagcagaggcgcgagctgctcttcaacgaggtggtaatcatgagggactaccagcacgag<br>aatgtggtggagatgtacaacagctacctggtgggggacgagctctgggtggtcatggagttcctggaag<br>gaggcgccctcaccgacatcgtcacccacaccaggatgaacgaggagcagatcgcagccgtgtgcctt<br>gcagtgctgcaggccctgtcggtgctccacgcccagggcgtcatccaccgggacatcaagagcgactcg<br>atcctgctgacccatgatggcagggtgaagctgtcagactttgggttctgcgcccaggtgagcaaggaagt<br>gccccgaaggaagtcgctggtcggcacgccctactggatggccccagagctcatctcccgccttccctac<br>gggccagaggtagacatctggtcgctggggataatggtgattgagatggtggacggagagcccccctact<br>tcaacgagccacccctcaaagccatgaagatgattcgggacaacctgccaccccgactgaagaacctg<br>cacaaggtgtcgccatccctgaagggcttcctggaccgcctgctggtgcgagaccctgcccagcgggcc<br>acggcagccgagctgctgaagcacccattcctggccaaggcagggccgcctgccagcatcgtgcccctc<br>atgcgccagaaccgcaccaga |
| pAG1910 | FIREtag-SGGGGSGG-mCherry-SGGS-INKAbox-PAK4cat | atgggtgacagatattgggtctttgtgaaacgggtgagcggcgggggaggctccggagggatggtgagc<br>aagggcgaggaggataacatggccatcatcaaggagttcatgcgcttcaaggtgcacatggagggctcc<br>gtgaacggccacgagttcgagatcgagggcgagggcgagggccgcccctacgagggcacccagacc<br>gccaagctgaaggtgaccaagggtggccccctgcccttcgcctgggacatcctgtcccctcagttcatgta<br>cggctccaaggcctacgtgaagcaccccgccgacatccccgactacttgaagctgtccttccccgagggc<br>ttcaagtgggagcgcgtgatgaacttcgaggacggcggcgtggtgaccgtgacccaggactcctccctgc<br>aggacggcgagttcatctacaaggtgaagctgcgcggcaccaacttcccctccgacggccccgtaatgc<br>agaagaagaccatgggctgggaggcctcctccgagcggatgtaccccgaggacggcgccctgaaggg<br>cgagatcaagcagaggctgaagctgaaggacggcggccactacgacgctgaggtcaagaccacctac<br>aaggccaagaagcccgtgcagctgcccggcgcctacaacgtcaacatcaagttggacatcacctccca<br>caacgaggactacaccatcgtggaacagtacgaacgcgccgagggccgccactccaccggcggcatg<br>gacgagctgtacaagtcaggtggatccgaagcagaggactggacagcagccctactgaacaggggtc<br>gcagtcgccagcccctggtactaggggacaattgctttgctgacttggtgcacaactggatggagctgcctg<br>aggaattccccgccgcccctgctgttcctgggcccctggcccccgctcaccacagcgggagccacagc<br>gagtatcccatgagcagttccgggctgccctgcagctggtggtggacccaggcgacccccgctcctacct<br>ggacaacttcatcaagattggcgagggctccacgggcatcgtgtgcatcgccaccgtgcgcagctcgggc |

| | | |
|---|---|---|
| | | aagctggtggccgtcaagaagatggacctgcgcaagcagcagaggcgcgagctgctcttcaacgaggt<br>ggtaatcatgagggactaccagcacgagaatgtggtggagatgtacaacagctacctggtgggggacga<br>gctctgggtggtcatggagttcctggaaggaggcgccctcaccgacatcgtcacccacaccaggatgaac<br>gaggagcagatcgcagccgtgtgccttgcagtgctgcaggccctgtcggtgctccacgcccagggcgtca<br>tccaccgggacatcaagagcgactcgatcctgctgacccatgatggcagggtgaagctgtcagactttgg<br>gttctgcgcccaggtgagcaaggaagtgccccgaaggaagtcgctggtcggcacgccctactggatggc<br>cccagagctcatctcccgccttccctacggccagaggtagacatctggtcgctggggataatggtgattg<br>agatggtggacggagagccccctacttcaacgagccacccctcaaagccatgaagatgattcgggac<br>aacctgccaccccgactgaagaacctgcacaaggtgtcgccatccctgaagggcttcctggaccgcctgc<br>tggtgcgagaccctgcccagcgggccacggcagccgagctgctgaagcacccattcctggccaaggca<br>gggccgcctgccagcatcgtgcccctcatgcgccagaaccgcaccaga |
| pAG1911 | $^{FIRE}$tag-SGGGGSGG-emiRFP670-SGGS-INKAbox-PAK4cat | atgggtgacagatattgggtctttgtgaaacgggtgagcggcgggggaggctccggagggatggcggaa<br>ggatccgtcgccaggcagcctgacctcttgacctgcgaacatgaagagatccacctcgccggctcgatcc<br>agccgcatggcgcgcttctggtcgtcagcgaacatgatcatcgcgtcatccaggccagcgccaacgccg<br>cggaatttctgaatctcggaagcgtactcggcgttccgctcgccgagatcgacggcgatctgttgatcaaga<br>tcctgccgcatctcgatcccaccgccgaaggcatgccggtcgcggtgcgctgccggatcggcaatccctct<br>acggagtactgcggtctgatgcatcggcctccggaaggcgggctgatcatcgaactcgaacgtgccggc<br>ccgtcgatcgatctgtcaggcacgctggcgccggcgctggagcggatccgcacggcgggttcactgcgc<br>gcgctgtgcgatgacaccgtgctgctgtttcagcagtgcaccggctacgaccgggtgatggtgtatcgtttcg<br>atgagcaaggccacggcctggtattctccgagtgccatgtgcctgggctcgaatcctatttcggcaaccgct<br>atccgtcgtcgactgtcccgcagatggcgcggcagctgtacgtgcggcagcgcgtccgcgtgctggtcga<br>cgtcacctatcagccggtgccgctggagccgcggctgtcgccgctgaccgggcgcgatctcgacatgtcg<br>ggctgcttcctgcgctcgatgtcgccgtgccatctgcagttcctgaaggacatgggcgtgcgcgccaccctg<br>gcggtgtcgctggtggtcggcggcaagctgtggggcctggttgtctgtcaccattatctgccgcgcttcatcc<br>gtttcgagctgcgggcgatctgcaaacggctcgccgaaaggatcgcgacgcggatcaccgcgcttgaga<br>gctcaggtggatccgaagcagaggactggacagcagccctactgaacaggggtcgcagtcgccagcc<br>cctggtactaggggacaattgctttgctgacttggtgcacaactggatggagctgcctgaggaattcccgc<br>cgcccctgctgttcctgggcccccctggcccccgctcaccacagcgggagccacagcgagtatcccatga<br>gcagttccgggctgccctgcagctggtggtggacccaggcgacccccgctcctacctggacaacttcatca<br>agattggcgagggctccacgggcatcgtgtgcatcgccaccgtgcgcagctcgggcaagctggtggccg<br>tcaagaagatggacctgcgcaagcagcagaggcgcgagctgctcttcaacgaggtggtaatcatgagg<br>gactaccagcacgagaatgtggtggagatgtacaacagctacctggtgggggacgagctctgggtggtc<br>atggagttcctggaaggaggcgccctcaccgacatcgtcacccacaccaggatgaacgaggagcagat<br>cgcagccgtgtgccttgcagtgctgcaggccctgtcggtgctccacgcccagggcgtcatccaccgggac<br>atcaagagcgactcgatcctgctgacccatgatggcagggtgaagctgtcagactttgggttctgcgccca<br>ggtgagcaaggaagtgccccgaaggaagtcgctggtcggcacgccctactggatggccccagagctca<br>tctcccgccttccctacggccagaggtagacatctggtcgctggggataatggtgattgagatggtggacg<br>gagagccccctacttcaacgagccacccctcaaagccatgaagatgattcgggacaacctgccacccc<br>gactgaagaacctgcacaaggtgtcgccatccctgaagggcttcctggaccgcctgctggtgcgagaccc<br>tgcccagcgggccacggcagccgagctgctgaagcacccattcctggccaaggcagggccgcctgcca<br>gcatcgtgcccctcatgcgccagaaccgcaccaga |
| pAG2054 | mAID-AS-EGFP-SGGS-INKAbox-PAK4cat | atgaaggagaagagtgcttgtcctaaagatccagccaaacctccggccaaggcacaagttgtgggatgg<br>ccaccggtgagatcataccggaagaacgtgatggtttcctgccaaaaatcaagcggtggcccggaggcg<br>gcggcgttcgtgaaggtatcaatggacggagcaccgtacttgaggaaaatcgatttgaggatgtataaag<br>ctagcatgagtaaaggagaagaacttttcactggagttgtcccaattcttgttgaattagatggtgatgttaatg<br>ggtacaaattttctgtcagtggagagggtgaaggtgatgcaacatacggaaaacttacccttaaatttatttg<br>cactactggaaaactacctgttccatggccaacacttgtcactactctcacttatggtgttcaatgcttttcaag<br>atatccagatcatatgaagcggcacgacttcttcaagagcgccatgcctgagggatacgtgcaggagag<br>gaccatcttcttcaaggacgacgggaactacaagacacgtgctgaagtcaagtttgagggagacaccctc<br>gtcaacaggatcgagcttaagggaatcgatttcaaggaggacggaaacatcctcggccacaagttggaa<br>tacaactacaactcccacaacgtatacatcatggccgacaagcaaaagaacggcatcaaagccaacttc<br>aagacccgccacaacatcgaagacggcggcgtgcaactcgctgatcattatcaacaaaatactccaatt<br>ggcgatgaccctgtccttttaccagacaaccattacctgtccacacaatctgccctttcgaaagatcccaac<br>gaaaagagagaccacatggtcattcttgagtttgtaacggctgctgggattacacatggcatggatgaacta<br>tacaaatcaggtggatccgaagcagaggactggacagcagccctactgaacaggggtcgcagtcgcca<br>gcccctggtactaggggacaattgctttgctgacttggtgcacaactggatggagctgcctgaggaattccc |

| | | |
|---|---|---|
| | | cgccgcccctgctgttcctgggccccctggcccccgctcaccacagcgggagccacagcgagtatcccat<br>gagcagttccgggctgccctgcagctggtggtggacccaggcgacccccgctcctacctggacaacttca<br>tcaagattggcgagggctccacgggcatcgtgtgcatcgccaccgtgcgcagctcgggcaagctggtgg<br>ccgtcaagaagatggacctgcgcaagcagcagaggcgcgagctgctcttcaacgaggtggtaatcatga<br>gggactaccagcacgagaatgtggtggagatgtacaacagctacctggtgggggacgagctctgggtgg<br>tcatggagttcctggaaggaggcgccctcaccgacatcgtcacccacaccaggatgaacgaggagcag<br>atcgcagccgtgtgccttgcagtgctgcaggccctgtcggtgctccacgcccagggcgtcatccaccggg<br>acatcaagagcgactcgatcctgctgacccatgatggcagggtgaagctgtcagactttgggttctgcgcc<br>caggtgagcaaggaagtgccccgaaggaagtcgctggtcggcacgccctactggatggccccagagct<br>catctcccgccttccctacgggccagaggtagacatctggtcgctggggataatggtgattgagatggtgga<br>cggagagccccccctacttcaacgagccacccctcaaagccatgaagatgattcgggacaacctgccacc<br>ccgactgaagaacctgcacaaggtgtcgccatccctgaagggcttcctggaccgcctgctggtgcgagac<br>cctgcccagcgggccacggcagccgagctgctgaagcacccattcctggccaaggcagggccgcctgc<br>cagcatcgtgcccctcatgcgccagaaccgcaccaga |
| pAG2055 | mAID-AS-<br>mCherry-<br>SGGS-<br>INKAbox-<br>PAK4cat | atgaaggagaagagtgcttgtcctaaagatccagccaaacctccggccaaggcacaagttgtgggatgg<br>ccaccggtgagatcataccggaagaacgtgatggtttcctgccaaaaatcaagcggtggcccggaggcg<br>gcggcgttcgtgaaggtatcaatggacggagcaccgtacttgaggaaaatcgatttgaggatgtataaag<br>ctagcatggtgagcaagggcgaggaggataacatggccatcatcaaggagttcatgcgcttcaaggtgc<br>acatggagggctccgtgaacggccacgagttcgagatcgagggcgagggcgagggccgcccctacga<br>gggcacccagaccgccaagctgaaggtgaccaagggtggccccctgcccttcgcctgggacatcctgtc<br>ccctcagttcatgtacggctccaaggcctacgtgaagcaccccgccgacatccccgactacttgaagctgt<br>ccttccccgagggcttcaagtgggagcgcgtgatgaacttcgaggacggcggcgtggtgaccgtgaccc<br>aggactcctccctgcaggacggcgagttcatctacaaggtgaagctgcgcggcaccaacttcccctccga<br>cggccccgtaatgcagaagaagaccatgggctgggaggcctcctccgagcggatgtaccccgaggac<br>ggcgccctgaagggcgagatcaagcagaggctgaagctgaaggacggcggccactacgacgctgag<br>gtcaagaccacctacaaggccaagaagcccgtgcagctgcccggcgcctacaacgtcaacatcaagtt<br>ggacatcacctcccacaacgaggactacaccatcgtggaacagtacgaacgcgccgagggccgccac<br>tccaccggcggcatggacgagctgtacaagtcaggtggatccgaagcagaggactggacagcagccct<br>actgaacaggggtcgcagtcgccagcccctggtactaggggacaattgctttgctgacttggtgcacaact<br>ggatggagctgcctgaggaattccccgccgcccctgctgttcctgggccccctggcccccgctcaccaca<br>gcgggagccacagcgagtatcccatgagcagttccgggctgccctgcagctggtggtggacccaggcg<br>acccccgctcctacctggacaacttcatcaagattggcgagggctccacgggcatcgtgtgcatcgccacc<br>gtgcgcagctcgggcaagctggtggccgtcaagaagatggacctgcgcaagcagcagaggcgcgagc<br>tgctcttcaacgaggtggtaatcatgagggactaccagcacgagaatgtggtggagatgtacaacagcta<br>cctggtgggggacgagctctgggtggtcatggagttcctggaaggaggcgccctcaccgacatcgtcacc<br>cacaccaggatgaacgaggagcagatcgcagccgtgtgccttgcagtgctgcaggccctgtcggtgctcc<br>acgcccagggcgtcatccaccgggacatcaagagcgactcgatcctgctgacccatgatggcagggtga<br>agctgtcagactttgggttctgcgcccaggtgagcaaggaagtgccccgaaggaagtcgctggtcggcac<br>gccctactggatggccccagagctcatctcccgccttccctacgggccagaggtagacatctggtcgctgg<br>ggataatggtgattgagatggtggacggagagccccccctacttcaacgagccacccctcaaagccatga<br>agatgattcgggacaacctgccaccccgactgaagaacctgcacaaggtgtcgccatccctgaagggctt<br>cctggaccgcctgctggtgcgagaccctgcccagcgggccacggcagccgagctgctgaagcacccatt<br>cctggccaaggcagggccgcctgccagcatcgtgcccctcatgcgccagaaccgcaccaga |
| pAG2062 | OsTIR1-<br>cMyc | atgacgtacttcccggaggaggtggtggagcacatcttcagcttcctgccggcgcagcgcgaccgcaaca<br>cggtctcgctcgtctgcaaggtgtggtacgagatcgagaggctgagccgccgcggcgtcttcgtgggcaa<br>ctgctacgccgtgcgcgccggccgcgtcgccgcgcggttccccaacgtgcgggcgctcacggtgaaggg<br>gaagccccacttcgccgacttcaacctcgtgcccccgactggggcggctacgcggggccgtggatcga<br>ggcggccgcgaggggatgccacggcctggaggagctcaggatgaagcggatggtggtgtccgacgag<br>agcctcgagctgctggctcgctcgttcccgcggttcagggctcttgttcttatcagctgcgaggggttcagcac<br>tgacgggctagccgccgtcgcgagccattgcaagcttctgagggagttggatttgcaggaaaatgaagtg<br>gaggatcgagggcctaggtggctttcctgcttccctgattcctgcacatcacttgtctcattgaattttgcctgca<br>tcaaaggggaggttaatgctggttcactggagagacttgttagcaggtccccaaacctgcggagtttgagg<br>ctgaatcgatctgtatcggtagatacacttgcaaagatactactgcgtaccccctaacttggaggatttggga<br>cagggaatttgacagatgacttccaaactgagtcctactttaagcttaccagtgctctggagaaatgcaaga<br>tgttgaggagtttgtctggattctgggatgcttctcctgtttgcctgtcatttatctacccccctgtgtgctcaactgac<br>aggattgaacttgagctatgcacccacacttgatgcttctgaccttacaaaaatgattagccgctgtgtgaag |

| | | |
|---|---|---|
| | | ctccaacgcctttgggtactggattgtatctcggacaaaggcttgcaagtggtggcctccagttgcaaagact<br>tgcaagaactcagggtatttccatcagatttctacgtagctggttattctgcagtgacagaggagggacttgtt<br>gcagtatccttgggctgtccaaaactgaactcactactgtacttctgtcaccaaatgactaatgctgcactagt<br>tactgtcgccaagaactgtccaaatttcacacgattcagactttgtattcttgagccagggaagcctgatgttgt<br>gacaagccaaccattagatgaaggctttggagctattgttcgtgagtgcaagggattacaacgtttgtcaata<br>tctggtcttctcacagacaaagttttcatgtatattgggaaatatgcaaaacaacttgagatgctttctatagcat<br>ttgctggtgacagtgataagggtatgatgcatgttatgaatggatgcaagaatttaaggaaactggagataa<br>gagatagcccgtttggtgatgctgcactcttggggaattttgctaggtacgagacaatgcgatccctttggatg<br>tcatcttgcaatgtcacgttaaaggggtgccaagtccttgcgtcaaagatgccgatgctcaatgttgaggtca<br>taaatgagcgggatggtagcaatgaaatggaggaaaaccatggagatctgcctaaagtggagaaattat<br>atgtgtaccgcacaactgctggggcgagggatgatgcaccaaattttgttaaaatcctaggatccgaacaa<br>aagcttatttctgaagaggacttg |
| pAG2063 | $OsTIR1_{F74G}$ - cMyc | atgacgtacttcccggaggaggtggtggagcacatcttcagcttcctgccggcgcagcgcgaccgcaaca<br>cggtctcgctcgtctgcaaggtgtggtacgagatcgagaggctgagccgccgcggcgtcttcgtgggcaa<br>ctgctacgccgtgcgcgccggccgcgtcgccgcgcggttccccaacgtgcgggcgctcacggtgaaggg<br>gaagccccacggcgccgacttcaacctcgtgcccccgactggggcggctacgcggggccgtggatcg<br>aggcggccgcgaggggatgccacggcctggaggagctcaggatgaagcggatggtggtgtccgacga<br>gagcctcgagctgctggctcgctcgttcccgcggttcagggctcttgttcttatcagctgcgaggggttcagca<br>ctgacgggctagccgccgtcgcgagccattgcaagcttctgagggagttggatttgcaggaaaatgaagt<br>ggaggatcgagggcctaggtggctttcctgcttccctgattcctgcacatcacttgtctcattgaattttgcctgc<br>atcaaaggggaggttaatgctggttcactggagagacttgttagcaggtccccaaacctgcggagtttgag<br>gctgaatcgatctgtatcggtagatacacttgcaaagatactactgcgtaccccctaacttggaggatttgggg<br>acagggaatttgacagatgacttccaaactgagtcctactttaagcttaccagtgctctggagaaatgcaag<br>atgttgaggagtttgtctggattctgggatgcttctcctgtttgcctgtcatttatctacccccctgtgtgctcaactga<br>caggattgaacttgagctatgcacccacacttgatgcttctgaccttacaaaaatgattagccgctgtgtgaa<br>gctccaacgcctttgggtactggattgtatctcggacaaaggcttgcaagtggtggcctccagttgcaaaga<br>cttgcaagaactcagggtatttccatcagatttctacgtagctggttattctgcagtgacagaggagggacttg<br>ttgcagtatccttgggctgtccaaaactgaactcactactgtacttctgtcaccaaatgactaatgctgcacta<br>gttactgtcgccaagaactgtccaaatttcacacgattcagactttgtattcttgagccagggaagcctgatgtt<br>gtgacaagccaaccattagatgaaggctttggagctattgttcgtgagtgcaagggattacaacgtttgtcaa<br>tatctggtcttctcacagacaaagttttcatgtatattgggaaatatgcaaaacaacttgagatgctttctatagc<br>atttgctggtgacagtgataagggtatgatgcatgttatgaatggatgcaagaatttaaggaaactggagat<br>aagagatagcccgtttggtgatgctgcactcttggggaattttgctaggtacgagacaatgcgatccctttgg<br>atgtcatcttgcaatgtcacgttaaaggggtgccaagtccttgcgtcaaagatgccgatgctcaatgttgagg<br>tcataaatgagcgggatggtagcaatgaaatggaggaaaaccatggagatctgcctaaagtggagaaat<br>tatatgtgtaccgcacaactgctggggcgagggatgatgcaccaaattttgttaaaatcctaggatccgaac<br>aaaagcttatttctgaagaggacttg |
| pAG2064 | rtTA3-cMyc | atgtctagactggacaagagcaaagtcataaacggagctctggaattactcaatggtgtcggtatcgaagg<br>cctgacgacaaggaaactcgctcaaaagctgggagttgagcagcctaccctgtactggcacgtgaagaa<br>caagcgggccctgctcgatgccctgccaatcgagatgctggacaggcatcatacccacttctgccccctgg<br>aaggcgagtcatggcaagactttctgcggaacaacgccaagtcataccgctgtgctctcctctcacatcgc<br>gacggggctaaagtgcatctcggcacccgcccaacagagaaacagtacgaaaccctggaaaatcagc<br>tcgcgttcctgtgtcagcaaggcttctccctggagaacgcactgtacgctctgtccgccgtgggccactttac<br>actgggctgcgtattggaggaacaggagcatcaagtagcaaaagaggaaagagagacacctaccacc<br>gattctatgcccccacttctgagacaagcaattgagctgttcgaccggcagggagccgaacctgccttcctt<br>tcggcctggaactaatcatatgtggcctggagaaacagctaaagtgcgaaagcggcgggccgaccgac<br>gcccttgacgattttgacttagacatgctcccagccgatgcccttgacgactttgaccttgatatgctgcctgct<br>ggatccgaacaaaagcttatttctgaagaggacttg |
| pAG2099 | DHB-linker-mVenus-SGGS-INKAbox-PAK4cat | atgacaaatgatgtcacctggagcgaggcctcttcgcctgatgagaggacactcacctttgctgaaagatg<br>gcaattatcttcacctgatggagtagatacagatgatgatttaccaaaatcgcgagcatccaaaagaacct<br>gtggtgtgaatgatgatgaaagtccaagcaaaatttttatggtgggagaatctccacaagtgtcttccagact<br>tcagaatttgagactgaataatttaattcccaggcaacttttcaagcccaccgataatcaagaaactggatcc<br>ggggcccagggcagcggcgtgagcaagggcgaggagctgttcaccggggtggtgcccatcctggtcga<br>gctggacggcgacgtaaacggccacaagttcagcgtgtccggcgagggcgagggcgatgccacctac<br>ggcaagctgaccctgaagctgatctgcaccaccggcaagctgcccgtgccctggcccaccctcgtgacc<br>accctgggctacggcctgcagtgcttcgcccgctaccccgaccacatgaagcagcacgacttcttcaagtc |

| | | |
|---|---|---|
| | | cgccatgcccgaaggctacgtccaggagcgcaccatcttcttcaaggacgacggcaactacaagaccc<br>gcgccgaggtgaagttcgagggcgacaccctggtgaaccgcatcgagctgaagggcatcgacttcaag<br>gaggacggcaacatcctggggcacaagctggagtacaactacaacagccacaacgtctatatcaccgc<br>cgacaagcagaagaacggcatcaaggccaacttcaagatccgccacaacatcgaggacggcggcgt<br>gcagctcgccgaccactaccagcagaacacccccatcggcgacggccccgtgctgctgcccgacaacc<br>actacctgagctaccagtccaagctgagcaaagaccccaacgagaagcgcgatcacatggtcctgctgg<br>agttcgtgaccgccgccgggatcactctcggcatggacgagctgtacaagtcaggtggatccgaagcag<br>aggactggacagcagccctactgaacaggggtcgcagtcgccagcccctggtactaggggacaattgct<br>ttgctgacttggtgcacaactggatggagctgcctgaggaattccccgccgcccctgctgttcctggcccc<br>ctggcccccgctcaccacagcgggagccacagcgagtatcccatgagcagttccgggctgccctgcagc<br>tggtggtggacccaggcgacccccgctcctacctggacaacttcatcaagattggcgagggctccacggg<br>catcgtgtgcatcgccaccgtgcgcagctcgggcaagctggtggccgtcaagaagatggacctgcgcaa<br>gcagcagaggcgcgagctgctcttcaacgaggtggtaatcatgagggactaccagcacgagaatgtggt<br>ggagatgtacaacagctacctggtgggggacgagctctgggtggtcatggagttcctggaaggaggcgc<br>cctcaccgacatcgtcacccacaccaggatgaacgaggagcagatcgcagccgtgtgccttgcagtgct<br>gcaggccctgtcggtgctccacgcccagggcgtcatccaccgggacatcaagagcgactcgatcctgct<br>gacccatgatggcagggtgaagctgtcagactttgggttctgcgcccaggtgagcaaggaagtgccccga<br>aggaagtcgctggtcggcacgccctactggatggccccagagctcatctcccgccttccctacgggccag<br>aggtagacatctggtcgctggggataatggtgattgagatggtggacggagagcccccctacttcaacga<br>gccacccctcaaagccatgaagatgattcgggacaacctgccaccccgactgaagaacctgcacaagg<br>tgtcgccatccctgaagggcttcctggaccgcctgctggtgcgagaccctgcccagcgggccacggcagc<br>cgagctgctgaagcacccattcctggccaaggcagggccgcctgccagcatcgtgcccctcatgcgcca<br>gaaccgcaccaga |
| pAG2101 | mCherry-linker-hGem(1-60)- SGGS-INKAbox-PAK4cat | atggtgagcaagggcgaggaggataacatggccatcatcaaggagttcatgcgcttcaaggtgcacatg<br>gagggctccgtgaacggccacgagttcgagatcgagggcgagggcgagggccgcccctacgagggc<br>acccagaccgccaagctgaaggtgaccaagggtggccccctgcccttcgcctgggacatcctgtcccctc<br>agttcatgtacggctccaaggcctacgtgaagcaccccgccgacatccccgactacttgaagctgtccttcc<br>ccgagggcttcaagtgggagcgcgtgatgaacttcgaggacggcggcgtggtgaccgtgacccaggact<br>cctccctgcaggacggcgagttcatctacaaggtgaagctgcgcggcaccaacttcccctccgacggccc<br>cgtaatgcagaagaagaccatgggctgggaggcctcctccgagcggatgtaccccgaggacggcgcc<br>ctgaagggcgagatcaagcagaggctgaagctgaaggacggcggccactacgacgctgaggtcaag<br>accacctacaaggccaagaagcccgtgcagctgcccggcgcctacaacgtcaacatcaagttggacat<br>cacctcccacaacgaggactacaccatcgtggaacagtacgaacgcgccgagggccgccactccacc<br>ggcggcatggacgagctgtacaagggatatccatcacactggcggccgctcgagatgaatcccagtatg<br>aagcagaaacaagaagaaatcaaagagaatataaagaatagttctgtcccaagaagaactctgaagat<br>gattcagccttctgcatctggatctcttgttggaagagaaaatgagctgtccgcaggcttgtccaaaaggaa<br>acatcggaatgaccacttaacatcttcaggtggatccgaagcagaggactggacagcagccctactgaa<br>caggggtcgcagtcgccagcccctggtactaggggacaattgctttgctgacttggtgcacaactggatgg<br>agctgcctgaggaattccccgccgcccctgctgttcctggcccccctggcccccgctcaccacagcggga<br>gccacagcgagtatcccatgagcagttccgggctgccctgcagctggtggtggacccaggcgacccccg<br>ctcctacctggacaacttcatcaagattggcgagggctccacgggcatcgtgtgcatcgccaccgtgcgca<br>gctcgggcaagctggtggccgtcaagaagatggacctgcgcaagcagcagaggcgcgagctgctcttc<br>aacgaggtggtaatcatgagggactaccagcacgagaatgtggtggagatgtacaacagctacctggtg<br>ggggacgagctctgggtggtcatggagttcctggaaggaggcgccctcaccgacatcgtcacccacacc<br>aggatgaacgaggagcagatcgcagccgtgtgccttgcagtgctgcaggccctgtcggtgctccacgccc<br>agggcgtcatccaccgggacatcaagagcgactcgatcctgctgacccatgatggcagggtgaagctgt<br>cagactttgggttctgcgcccaggtgagcaaggaagtgccccgaaggaagtcgctggtcggcacgccct<br>actggatggccccagagctcatctcccgccttccctacgggccagaggtagacatctggtcgctggggata<br>atggtgattgagatggtggacggagagcccccctacttcaacgagccacccctcaaagccatgaagatg<br>attcgggacaacctgccaccccgactgaagaacctgcacaaggtgtcgccatccctgaagggcttcctgg<br>accgcctgctggtgcgagaccctgcccagcgggccacggcagccgagctgctgaagcacccattcctgg<br>ccaaggcagggccgcctgccagcatcgtgcccctcatgcgccagaaccgcaccaga |
